# Structure of zebrafish NLRP3 reveals a novel mode of inflammasome activation

**DOI:** 10.64898/2026.04.17.719140

**Authors:** Lisa S. Dopslaff, María Mateo-Tórtola, Varvara Varlamova, Corinna Gehring-Khav, Marit H. Walle, Lukas Schenk, Alexander N.R. Weber, Veit Hornung, Liudmila Andreeva

**Affiliations:** Department of Medical Oncology and Pneumology, University Hospital Tübingen, Tübingen, Germany; iFIT – Cluster of excellence (EXC 2180). “Image-Guided and Functionally Instructed Tumor Therapies”, University of Tübingen, Tübingen, Germany; Institute of Immunology, Department of Innate Immunity, University of Tübingen, Tübingen, Germany; Gene Center and Department of Biochemistry, Ludwig-Maximilians-Universität, Munich, Germany; CMFI – Cluster of excellence (EXC 2124). “Controlling Microbes to Fight Infections”, Tübingen, Germany; German Cancer Consortium (DKTK), DKTK Partner Site Tübingen, Tübingen, Germany

**Keywords:** NLRP3, inflammasome, zebrafish, inflammation, innate immunity, cryo-EM

## Abstract

NLRP3 is an innate immune sensor of a broad range of stimuli, which upon activation forms a multiprotein inflammasome complex triggering caspase-1 activation, IL-1β and IL-18 maturation, and inflammatory cell death. The canonical NLRP3 activation pathway has been well characterized from a structural perspective. It involves the association of NLRP3 with membranes in the form of inactive oligomeric “cage” complexes, which, upon activation, convert to an active oligomeric NLRP3 disc. NLRP3 structural rearrangements during non-classical NLRP3 activation pathways, however, remain unknown. Here, we report a novel mode of NLRP3 activation utilized by the NLRP3 homolog from zebrafish. The cryo-EM structure of zebrafish NLRP3 shows that, unlike human NLRP3, it forms disc-shaped heptamers that undergo further trimerization, resulting in a 21-mer oligomeric arrangement. Surprisingly, a single zebrafish NLRP3 heptamer cannot arrange its PYD domains into a PYD helix and therefore requires a trimer of heptamers to form a PYD filament that enables ASC oligomerization. Furthermore, zebrafish NLRP3 does not associate with the Golgi network, nor does it form inactive “cage” oligomers or interact with NEK7. Thus, our data demonstrate an ancestral non-canonical structural mechanism of NLRP3 activation, which may shed light on alternative NLRP3 activation pathways present in humans.

## Introduction

The NLRP3 inflammasome is an innate immune sensor that triggers cell death and inflammation in response to various infections and endogenous damage signals. It is involved in numerous inflammatory conditions including autoinflammatory cryopyrin-associated periodic syndromes (CAPS), cancer, and metabolic disorders, making NLRP3 a key regulator of inflammation and an attractive target for research and therapy development^1–5^.

In the ‘canonical’ NLRP3 activation pathway, inactive NLRP3 gets activated through a two-step process. The first “priming” step involves transcriptional upregulation of inflammasome components and post-translational modifications of NLRP3^6,7^. During the second step, NLRP3 gets activated by specific stimuli, which trigger NLRP3 oligomerization, recruitment of the adaptor protein apoptosis-associated speck-like protein containing a CARD (ASC), which in turn recruits the proinflammatory caspase-1. Formation of the inflammasome complex activates caspase-1 leading to maturation and release of IL-1β and IL-18, GSDMD cleavage and pyroptotic cell death^8–10^. Mechanistically, NLRP3-activating stimuli trigger the trafficking of the membrane- and Golgi-bound inactive NLRP3 “cage” oligomers^11–13^ to the microtubule organization center (MTOC)^11,14^, where, with the help of a co-factor, NIMA-related kinase 7 (NEK7), it gets disrupted and rearranged into an active 10-/11-fold NLRP3 disc, which initiates inflammasome assembly^15^. Finally, multiple inflammasomes and ASC oligomers assemble at the site of NLRP3 activation, forming an ASC speck^16,17^.

Apart from the canonical pathway, several non-classical strategies of NLRP3 activation have been reported. First, in human monocytes, a priming stimulus alone was sufficient to activate NLRP3 without ASC speck formation and cell death^18,19^. Similarly, lipopolysaccharide (LPS) stimulation alone could activate NLRP3 in mouse bone marrow-derived dendritic cells^20^ and human neutrophils^21,22^. Second, human^23^ and mouse^24^ myeloid cells under certain conditions were demonstrated to bypass the requirement for NEK7, which has been previously established as an essential NLRP3 co-factor^25–27^. Furthermore, non-Golgi-associated NLRP3 can also undergo activation and form ASC specks distally from the MTOC in parallel with canonical NLRP3 activation, albeit more slowly^28,29^. This MTOC-independent pathway did not require NLRP3 association with the Golgi nor oligomeric “cage” formation^28^ and could be recapitulated *in vitro* by mutation of palmitoylation sites^30^ or deletion of exon 3 in human *NLRP3*^28^, which encodes a flexible linker connecting pyrin (PYD) and fish-specific NACHT-associated (FISNA) domains and was shown to harbor one of the NLRP3 palmitoylation sites^30,31^. Such differential requirements for co-factors and localization sites suggests that additional, currently unexplored sets of structural ‘acrobatics’ may be at play in these non-classical NLRP3 activation pathways.

To address alternative mechanisms of NLRP3 activation that occur in nature we decided to elucidate structural rearrangements of NLRP3 in ancestral species. Indeed, the *NLRP3* gene was previously found not only in mammals, but also in birds, reptiles, amphibians and even teleost fishes^32^ (Fig. 1a). While the ability of NLRP3 to form an active inflammasome in lower vertebrates is poorly elucidated, NLRP3 was demonstrated to be functional in activation in teleosts such as Japanese flounder^33^, turbot^34^, yellow croaker^35^ and zebrafish^36,37^. Zebrafish NLRP3 consists of PYD, FISNA, NACHT, leucine-rich-repeat (LRR) and B30.2 domains (Fig. 1a), and was shown to interact with zebrafish ASC and caspase-1 homologs, caspases-A and -B, to trigger maturation of zebrafish IL-1β and GSDME cleavage leading to cell death^36^ (Supplementary Fig. 1a). We therefore decided to explore the structural aspects of zebrafish NLRP3 to gain insight into fundamental and potentially conserved features of NLRP3 activation in nature.

**Fig. 1.**
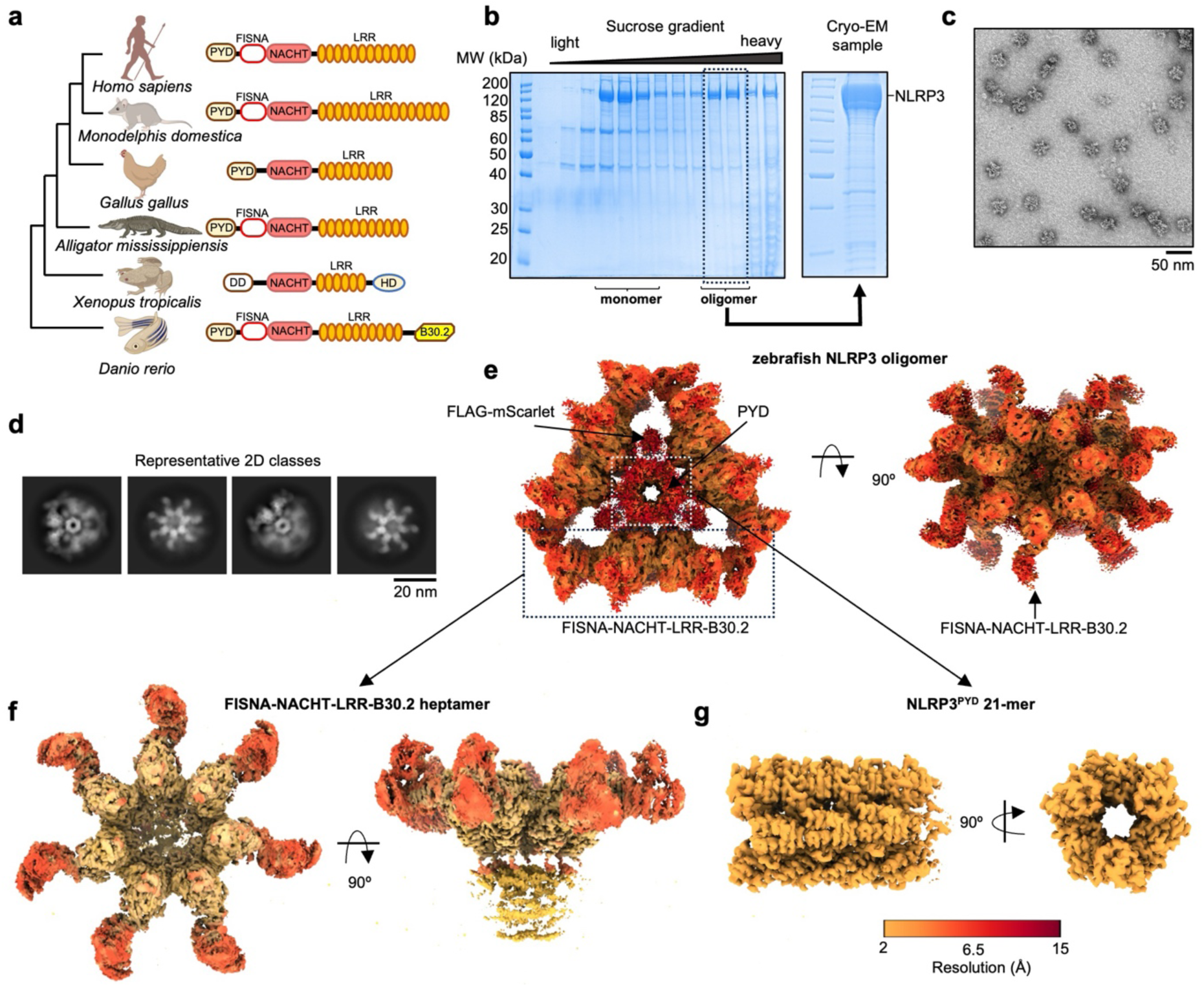
Purification and overall structure of zebrafish NLRP3. **a**, Domain structure of NLRP3 homologs from vertebrates. **b**, SDS-PAGE gel of sucrose gradient fractions of zebrafish NLRP3 (left), fractions collected for cryo-EM (dashed box) and SDS-PAGE gel of the final sample (right). **c**, Representative negative-staining EM image of zebrafish NLRP3 oligomer (scale bar 50 nm). **d,** Representative 2D class averages of zebrafish NLRP3 oligomers (scale bar 20 nm). **e-g,** Top and side views of the zebrafish NLRP3 oligomer cryo-EM maps: an initial cryo-EM map (**e**) and cryo-EM maps of the disc-shaped zebrafish NLRP3 heptamer (**f**) and PYD 21-mer (**g**) after focused refinement. The maps are colored by resolution.

In this study, we demonstrate a novel non-canonical mode of NLRP3 activation utilized by zebrafish NLRP3. The cryo-electron microscopy (cryo-EM) structure of the active zebrafish NLRP3 reveals formation of an unusual 21-mer oligomeric complex consisting of three disc-shaped NLRP3 heptamers, whose joint 21-mer PYD filament can trigger downstream signaling. Zebrafish NLRP3 formed an active inflammasome in human cells but did not rely on Golgi localization, “cage” formation or NEK7 binding. Thus, this ancestral NLRP3 activation pathway offers a previously unknown alternative mechanism for forming an active NLRP3 complex and may provide mechanistic insights about currently underexplored non-classical NLRP3 activation pathways in humans.

## Results

### Cryo-EM structure of zebrafish NLRP3 reveals an unusual oligomeric complex consisting of three disc-shaped NLRP3 heptamers

For structural studies, N-terminally FLAG- and mScarlet-tagged full-length zebrafish NLRP3 (NCBI MN088121.1) was expressed in Expi293F cells and purified using anti-FLAG affinity chromatography followed by ultracentrifugation on a sucrose gradient. Zebrafish NLRP3 was observed in both light (20-30%) and heavy (40-45%) sucrose gradient fractions (Fig. 1b) corresponding to its monomeric and oligomeric forms, respectively. Negative staining electron microscopy (EM) of zebrafish NLRP3 oligomers showed unexpected star-shaped complexes with a diameter of ∼ 30 nm (Fig. 1c), which were used for cryo-EM data acquisition. An initial cryo-EM map revealed a full zebrafish NLRP3 complex at 3.97 Å resolution (Fig. 1d, e, Supplementary Fig. 1b, c). In this complex, three disc-shaped FISNA-NACHT-LRR-B30.2 heptamers are rotated ∼ 60° to each other, forming a triangular structure with a PYD 21-mer in the middle (Fig. 1e). The density between the heptamers is likely emerging from the N-terminal FLAG-mScarlet tag, which was not removed during the purification.

Since the flexibility of the heptamers respective to each other obscured the resolution, we performed a focused refinement on different parts of the structure and obtained a 3.18 Å resolution map of a single zebrafish NLRP3 heptamer and a 2.62 Å resolution map of the PYD 21-mer (Fig. 1 f, g, Supplementary Fig. 1c, Supplementary Fig. 2a, b and Supplementary Table 1). Local resolution estimation indicated a lower resolution of the C-terminal LRR-B30.2 region (Fig. 1f), though all zebrafish NLRP3 domains were clearly visible. These cryo-EM maps were further used for model building and refinement.

### Zebrafish NLRP3 utilizes NACHT-NACHT, LRR-LRR and PYD-PYD interactions to stabilize a novel 21-mer conformation

The zebrafish NLRP3 oligomer consists of three disc-shaped FISNA-NACHT-LRR-B30.2 heptamers connected by a short 21-mer PYD filament inside of the complex, and by LRR-LRR interactions between adjacent discs forming a mostly charged 273.8 Å^2^ interface I (Fig. 2a-c, Supplementary Fig. 3a). Indeed, mutations within the interface I or deletion of the PYD abolished zebrafish NLRP3 oligomer formation (Fig. 2d, Supplementary Fig. 3b). Interestingly, stabilization of oligomeric complexes by PYD-PYD interactions has been also observed for the active human NLRP3 discs^15^ and inactive mouse NLRP3 “cages”^11^, indicating a similar role of NLRP3^PYD^ in both signal transduction and NLRP3 oligomer stabilization.

**Fig. 2.**
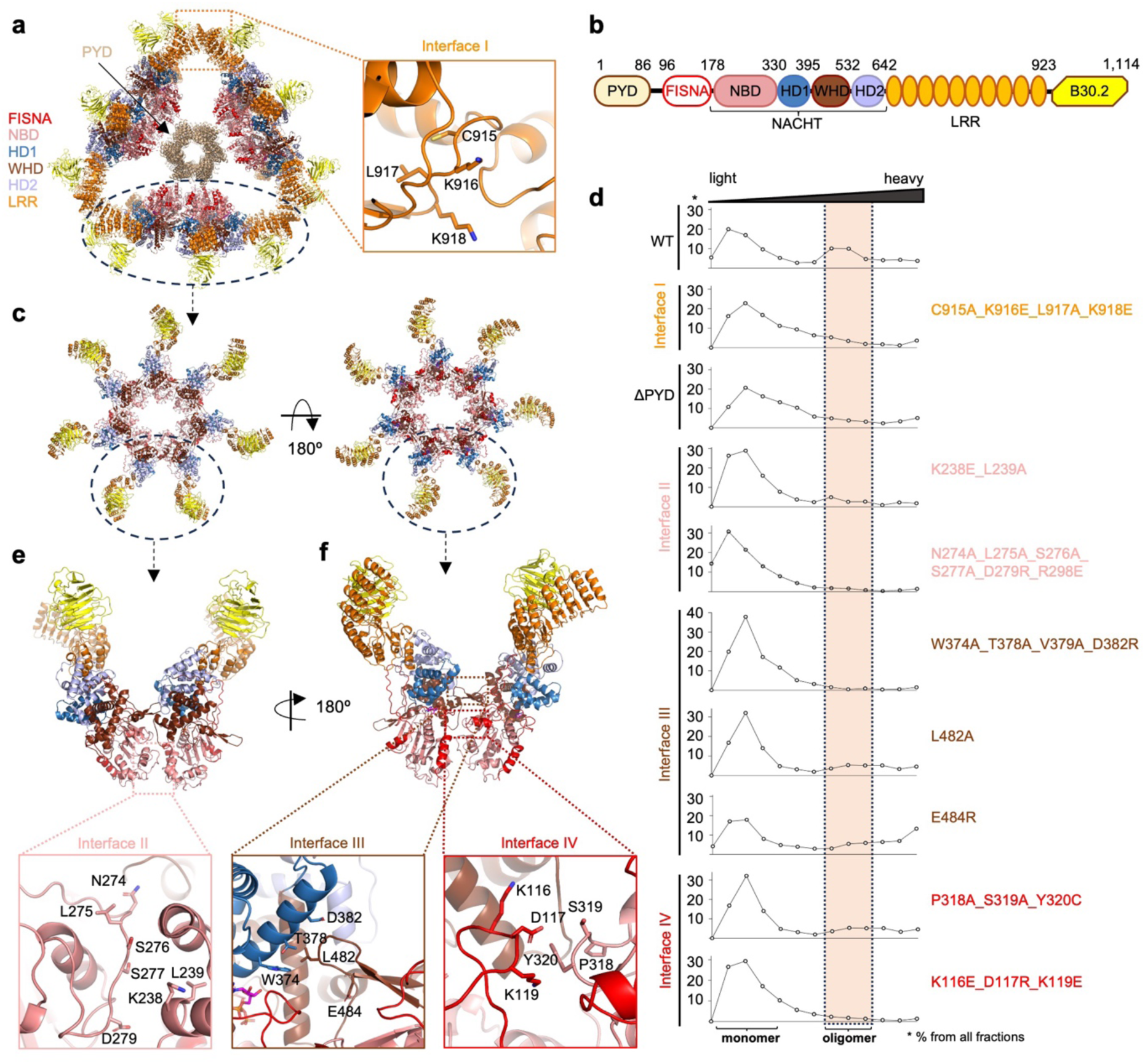
A model of the zebrafish NLRP3 oligomer and its main interfaces. **a**, Atomic model of the full zebrafish NLRP3 21-mer assembly, colored by domains, and a detailed view of the interface I. **b**, Domain structure of the zebrafish NLRP3. **c**, Top and bottom views of the atomic model of a disc-shaped zebrafish NLRP3 heptamer, colored by domains. **d**, Sucrose gradient profiles of wild-type (WT) and mutant zebrafish NLRP3 color-coded by interface: orange for interface I, salmon for interface II, brown for interface III and red for interface IV. The profiles were calculated by quantification of zebrafish NLRP3 bands from the SDS-PAGE images in Supplementary Fig. 3b. Highlighted are fractions containing NLRP3 oligomer. **e, f**, Overview and detailed views of interfaces II (**e**), III and IV(**f**) stabilizing the disc-shaped zebrafish NLRP3 heptamer, colored by domains. Residues used for mutagenesis are represented as sticks.

NACHT domains within a single disc-shaped zebrafish NLRP3 heptamer adopted an “open” conformation similar to the reported active oligomeric NLRC4 “wheel” structures^38–40^ and the active human NLRP3 disc structure^15^ (Supplementary Fig. 3c). Together with the ability to form a PYD 21-mer bundle (Fig. 1e), this suggests that the zebrafish 21-mer structure is an active NLRP3 complex. The 1,030.4 Å^2^ interface between NACHT domains of zebrafish NLRP3 is mostly charged (Supplementary Fig. 3d, e) and is stabilized by nucleotide-binding domain (NBD)-NBD (Interface II, Fig. 2e), winged helix domain (WHD) β-hairpin-helical domain 1 (HD1, Interface III, Fig. 2f) and NBD-FISNA (Interface IV, Fig. 2f) subdomain interactions. Mutation of one of these interfaces disrupted zebrafish NLRP3 oligomer formation (Fig. 2d, Supplementary Fig. 3b).

The overall zebrafish NLRP3 disc arrangement demonstrates several unique features as compared to human NLRP3. First, unlike 10 or 11 monomers required for the active human NLRP3 complex^15^, zebrafish NLRP3 requires only 7 NLRP3 monomers (Fig. 2c, Supplementary Fig. 4a), which so far has been observed only for Apaf-1 apoptosome^41–43^ but not for other NLRs and inflammasomes. Second, the LRR domains of zebrafish NLRP3 are oriented anticlockwise in contrast to their clockwise orientation in the active human NLRP3 disc (Fig. 2c, Supplementary Fig. 4a). Such an arrangement emerges from differences in helical domains 2 (HD2) between various teleost and tetrapod species, which are responsible for the orientation of the LRR domain (Supplementary Fig. 4b) but likely do not participate in protein-protein contacts in the active NLRP3 discs. Third, the concave LRR surfaces in the zebrafish NLRP3 heptamer are occupied by the C-terminal B30.2 domains instead of NEK7 as in the human active NLRP3 disc (Fig. 2c, Supplementary Fig. 4a). In sum, zebrafish NLRP3 utilizes interfaces similar to those stabilizing the active human NLRP3 complex to achieve a completely different oligomeric conformation. This highlights the plasticity of NLRP3 in forming oligomers of various shapes.

### A unique conformation of the FISNA domain stabilizes the active conformation of the NACHT domain of zebrafish NLRP3

Although the inactive conformation of zebrafish NLRP3 is unknown, we modeled it using AlphaFold 3^44^ based on the human NLRP3 inactive state^12^ in order to address possible structural rearrangements of zebrafish NLRP3 upon activation. Since a second FISNA helix, which was ordered the AlphaFold 3 predictions (“helix 2” in Supplementary Fig. 5a), was disordered in all available inactive human NLRP3 structures^12,13,45^, we modeled an inactive zebrafish NLRP3 with the second FISNA helix as disordered and the NACHT domain in a conformation identical to human inactive NLRP3^12^ (Fig. 3a, Supplementary Fig. 5b). Based on this model, upon activation zebrafish NLRP3 would undergo conformational changes in the NACHT domain similar to those in human NLRP3, in which the WHD-HD2-LRR-B30.2 segment rotates by 85.3° respective to the FISNA-NBD-HD1 domains (Fig. 3a).

**Fig. 3.**
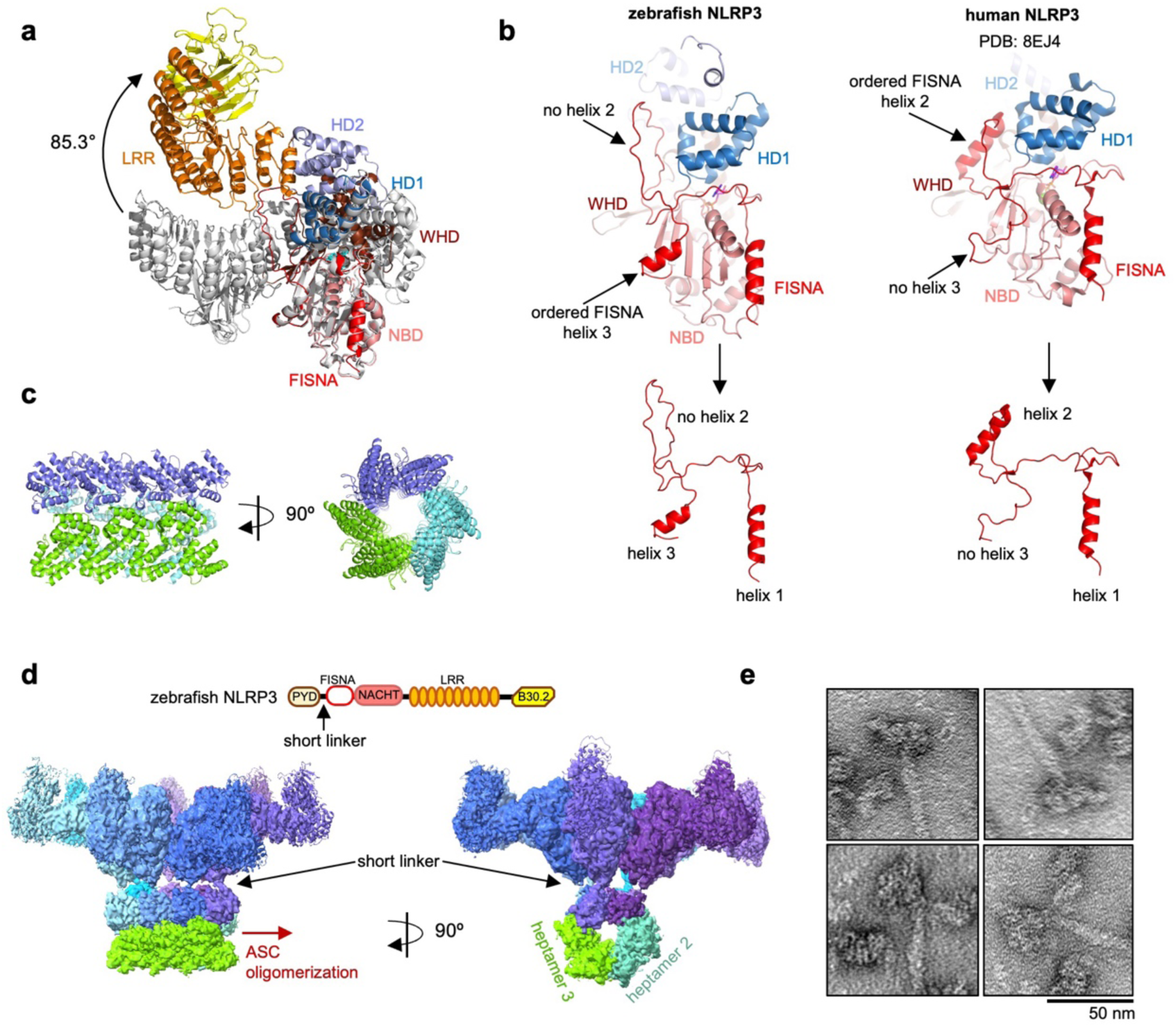
Conformational changes of zebrafish NLRP3 upon activation. **a**, Superposition of active zebrafish NLRP3 from the cryo-EM structure (colored by domain) and inactive zebrafish NLRP3 (grey) modeled based on inactive human NLRP3 (PDB: 7PZC). Direction of rotation of the WHD-HD2-LRR-B30.2 segment is indicated with an arrow. **b,** Comparison of FISNA domain position and structure in zebrafish (left) and human (right) NLRP3 demonstrating differences in conformation of the three key FISNA elements, colored by domains. **c**, Top and side views of the atomic model of the zebrafish NLRP3 PYD 21-mer, colored by the heptamer they belong to. **d**, Domain structure of zebrafish NLRP3 with the PYD-FISNA linker highlighted (top) and a fragment of the cryo-EM map from Fig. 1e showing a single disc-shaped heptamer connected to the PYD 21-mer, colored by monomer. PYD domains corresponding to the additional two 7-fold discs are colored cyan and green. The proposed direction of ASC oligomerization is indicated with a red arrow. **e**, Representative negative-staining EM images of zebrafish NLRP3 oligomer in complex with zebrafish ASC^PYD^ (scale bar 50 nm).

Like in human NLRP3, the “helix 2” FISNA region of the zebrafish NLRP3 (aa 134-162, in human aa 176-202) becomes reorganized upon activation, but does not form an α-helix present in the active form of human NLRP3^15^ (Fig. 3b, Supplementary Fig. 5c). Instead, the zebrafish FISNA forms an α-helix (“helix 3”) in a region adjacent to the NBD, which is absent in the active human NLRP3 structure (Fig. 3b, Supplementary Fig. 5c).

Remarkably, according to a PDBePISA^46^ macromolecular interface analysis, the “helix 2” region of zebrafish NLRP3 has an extensive 1,390.9 Å^2^ interface with all NACHT subdomains within an active NLRP3 monomer (Supplementary Fig. 5d), which is much larger than a similar 884.8 Å^2^ interface in human NLRP3 (Supplementary Fig. 5e). While human FISNA extensively participates in NLRP3 contacts in trans^15^ (Supplementary Fig. 5f), zebrafish FISNA domains are rotated further relative to each other (45.06° in 7-fold zebrafish vs. 36° in human 10-fold disc) (Supplementary Fig. 5g), which drastically reduces their interaction with adjacent NLRP3 monomers. Overall, it is likely that the zebrafish NLRP3 FISNA domain plays a stronger role in stabilizing the active conformation of an NLRP3 monomer than in human NLRP3.

### Trimerization of zebrafish NLRP3 disc-shaped heptamers is required for PYD nucleation and downstream signaling

The final atomic model of the zebrafish NLRP3 PYD 21-mer (Fig. 3c) contains 21 PYDs corresponding to three disc-shaped heptamers formed by FISNA-NACHT-LRR-B30.2 domains (Fig. 1e, Fig. 2a). Based on the symmetry search in CryoSparc, zebrafish NLRP3^PYD^ forms a right-handed helix with a 56° helical twist and 14.2 Å axial rise (Supplementary Fig. 1d), which surprisingly is nearly identical to the symmetry known for the human NLRP3^PYD^ filament^47^. Although the surface charge of the zebrafish NLRP3^PYD^ 21-mer is different from the human NLRP3^PYD 47^, it is still likely compatible with both zebrafish and human ASC (Supplementary Fig. 6a-c).

Interestingly, whereas the human NLRP3 nucleates PYD filaments *perpendicularly* to the disc plane (Supplementary Fig. 6d), the zebrafish PYD 21-mer is oriented *parallel* to the disc-shaped heptamer (Fig. 1e, Fig. 2a, Fig. 3d). Unlike human *NLRP3* gene, which encodes a 39 aa linker in exon 3 and enables flexible PYD positioning, this exon it is absent in zebrafish *NLRP3* (Supplementary Fig. 6e). Thus, a single NLRP3 heptamer cannot form a PYD filament by itself; instead, 7 PYDs from each heptamer cooperate to form a 21-mer complex with a full 21-mer PYD filament (Fig. 3d). In this arrangement none of the three heptameric FISNA-NACHT-LRR-B30.2 discs sterically hinders filament growth (Fig. 2a, Fig. 3d). Indeed, the zebrafish NLRP3 21-mer nucleated zebrafish ASC^PYD^ oligomerization (Fig. 3e). Collectively, these data clearly indicate that, despite an unexpected alternative geometry compared to human NLRP3, the 21-mer of zebrafish NLRP3 is active in ASC recruitment.

### ATP binding stabilizes but does not trigger formation of the active zebrafish NLRP3 oligomer

According to the current hypothesis, NLRP3 activation is accompanied by an exchange of ADP with ATP within the NBD domain^48–50^. This involves coordination by ATP-binding motifs, which are conserved in the STAND AAA+ ATPases superfamily to which NLRP3 belongs^51,52^. All residues involved in ATP binding and hydrolysis^49,53^ were conserved in both zebrafish and human NLRP3 (Fig. 4a), suggesting the importance of ATP binding for zebrafish NLRP3 activation. Indeed, a density corresponding to ATP was observed in all NLRP3 monomers within the active 21-mer zebrafish NLRP3 arrangement (Fig. 4b, Supplementary Fig. 7a). Of note, we did not include Mg^2+^ in the model, since no strong corresponding density was observed. Since active oligomeric zebrafish NLRP3 was purified from unstimulated cells in the presence of ATP, we were wondering whether ATP binding alone could have been sufficient for triggering the active zebrafish NLRP3 complex formation. To test this hypothesis, we purified zebrafish NLRP3 in the presence of ADP, ATP, the non-hydrolyzable ATP analog AMP-PNP, or without the addition of any nucleotide. We observed the formation of a similar proportion of an active oligomer, as judged by ultracentrifugation in a sucrose gradient (Fig. 4c, Supplementary Fig. 7b). Remarkably, the presence of AMP-PNP did not affect the 21-mer zebrafish NLRP3 peak but induced a shift of the monomeric fraction towards heavier intermediate species (Fig. 4c, Supplementary Fig. 7b). Negative staining EM of this sample revealed the presence of various multimeric states of zebrafish NLRP3 in the heavy sucrose gradient fractions (Fig. 4c). Taken together, these observations suggest that ATP binding can trigger some zebrafish NLRP3 multimerization but does not suffice for formation of the active complex. Thus, the active zebrafish NLRP3 oligomer must have been formed already during the expression.

**Fig. 4.**
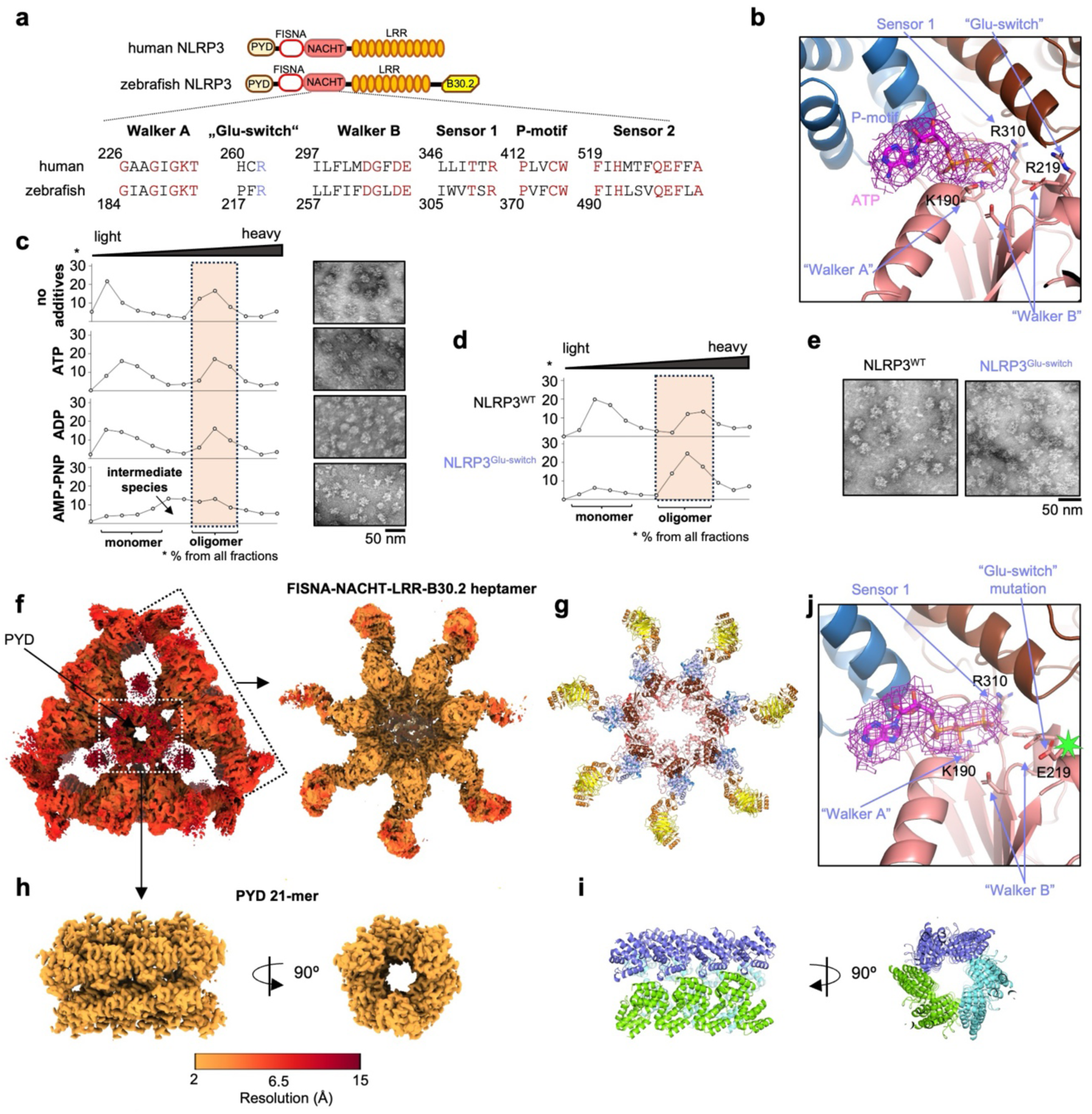
ATP binding and cryo-EM structure of the “Glu-switch”-mutated zebrafish NLRP3. **a**, Position and sequence of the main NLRP3 elements responsible for ATP binding and hydrolysis. Consensus residues with a high degree of conservation in all human NLRs are indicated in red. Arginine (R) 262 mutated in a gain-of-function version of human NLRP3 is conserved in zebrafish NLRP3 and is indicated in blue. **b**, A detailed view of the zebrafish NLRP3 ATP-binding pocket. Cryo-EM density of ATP is shown as isomesh at 1 root mean square deviation (RMSD) with a 2.5 Å radius. **c,** Sucrose gradient profiles and representative negative-staining EM images of WT zebrafish NLRP3 purified without or in presence of the indicated nucleotides (scale bar 50 nm). The profiles were calculated by quantification of zebrafish NLRP3 bands from the SDS-PAGE images in Supplementary Fig. 7b. Highlighted are fractions used for negative staining EM. **d, e**, Sucrose gradient profiles (**d**) and representative negative-staining EM images (**e**) of WT and “Glu-switch”-mutated zebrafish NLRP3 (scale bar 50 nm). The profiles were calculated by quantification of zebrafish NLRP3 bands from the SDS-PAGE images in Supplementary Fig. 7c. Highlighted are fractions containing NLRP3 oligomer. **f-i**, Cryo-EM maps (**f, h**) colored by resolution and atomic models (**g, i**) of the “Glu-switch”-mutated zebrafish NLRP3 oligomer. The atomic model of the disc-shaped NLRP3 heptamer (**g**) is colored by domains. Separate PYD domains in the atomic model of the PYD 21-mer (**i**) are colored by the heptamer they belong to. **j**, A detailed view of the “Glu-switch”-mutated zebrafish NLRP3 ATP-binding pocket. Cryo-EM density of ATP is shown as isomesh at 1 root mean square deviation (RMSD) with a 2.5 Å radius. The R219E mutation is labeled with a green star.

To further elucidate the involvement of ATP binding and hydrolysis in zebrafish NLRP3 activation, we mutated residues within the “Glu-switch” motif^54^, including R219, which corresponds to human R262 (Fig. 4a). In human NLRP3, mutation of R262 results in its constitutive activation and CAPS disease^55–57^. Remarkably, zebrafish NLRP3 with an analogous mutation in the “Glu-switch” region (R219E-E220R-L223A) formed nearly exclusively active oligomers, as opposite to only ∼30% for the wild-type protein (Fig. 4d, e, Supplementary Fig. 7c). Thus, the “Glu-switch” mutation of zebrafish NLRP3 is likely to result in its gain-of-function form.

Various gain-of-function mutations of human NLRP3 have been recently reported to have differential requirements for priming and activation steps^58^, raising the question of whether some NLRP3 disease variants might deviate from a canonical two-step mechanism of activation. Since no structure of NLRP3 with a gain-of-function mutation has been reported so far, we wanted to address possible structural changes induced by such mutations. For this purpose, we collected a cryo-EM dataset of the “Glu-switch”-mutated zebrafish NLRP3. However, a 4.16 Å initial cryo-EM map of the complex revealed the same arrangement as observed for a wild-type zebrafish NLRP3 (Fig. 4f, Supplementary Fig. 8, Supplementary Table 1). Using the focused refinement, we improved the FISNA-NACHT-LRR-B30.2 heptamer and PYD 21-mer densities to resolutions of 3.12 Å and 2.67 Å, respectively, and built an atomic model of the “Glu-switch”-mutated zebrafish NLRP3 (Fig. 4f-i, Supplementary Fig. 8d). “Glu-switch”-mutated zebrafish NLRP3 contained ATP bound to every NLRP3 monomer (Fig. 4j, Supplementary Fig. 7d) and was in general highly similar to the wild-type protein (RMSD < 1.4Å between heptamers). In human NLRP3, a comparable “Glu-switch” mutation reduced ATP hydrolysis and thus increased the retention time of the active ATP-bound NLRP3 conformation^49^, which explains why at least the R219E-E220R-L223A mutation in the “Glu-switch” motif did not result in a structurally different active NLRP3 oligomer.

### Zebrafish NLRP3 forms an active inflammasome in human cells

Since the active zebrafish NLRP3 21-mer can nucleate ASC oligomerization (Fig. 3e), we were wondering whether it would form active inflammasomes in human cells. For this purpose, inducible NLRP3 constructs were reconstituted into human monocyte-like BlaER1 NLRP3-KO cells^59^ and the inflammasome-dependent cell death was measured by release of lactate dehydrogenase (LDH). Whereas NLRP3 homologs from other teleosts were shown to respond to canonical NLRP3 stimuli like human NLRP3^33–35^, under the conditions tested, we were not able to detect a stimulus-dependent zebrafish NLRP3 activation. Instead, zebrafish NLRP3 demonstrated high spontaneous activity (Fig. 5a, Supplementary Fig. 9a). Substitution of the zebrafish PYD domain with the human PYD further increased constitutive activity of NLRP3, as judged by LDH release in BlaER1 cells (Fig. 5a, Supplementary Fig. 9a). Similar results were observed in monocyte-like THP-1 NLRP3-KO cells reconstituted with NLRP3 constructs, in which zebrafish NLRP3 constructs spontaneously initiated GSDMD cleavage (Fig. 5b). In agreement with the surface charge of the PYD filaments (Supplementary Fig. 6a-c), these results confirm that zebrafish NLRP3 is compatible with human ASC and triggers ASC speck formation, GSDMD cleavage and cell death in human cells.

**Fig. 5.**
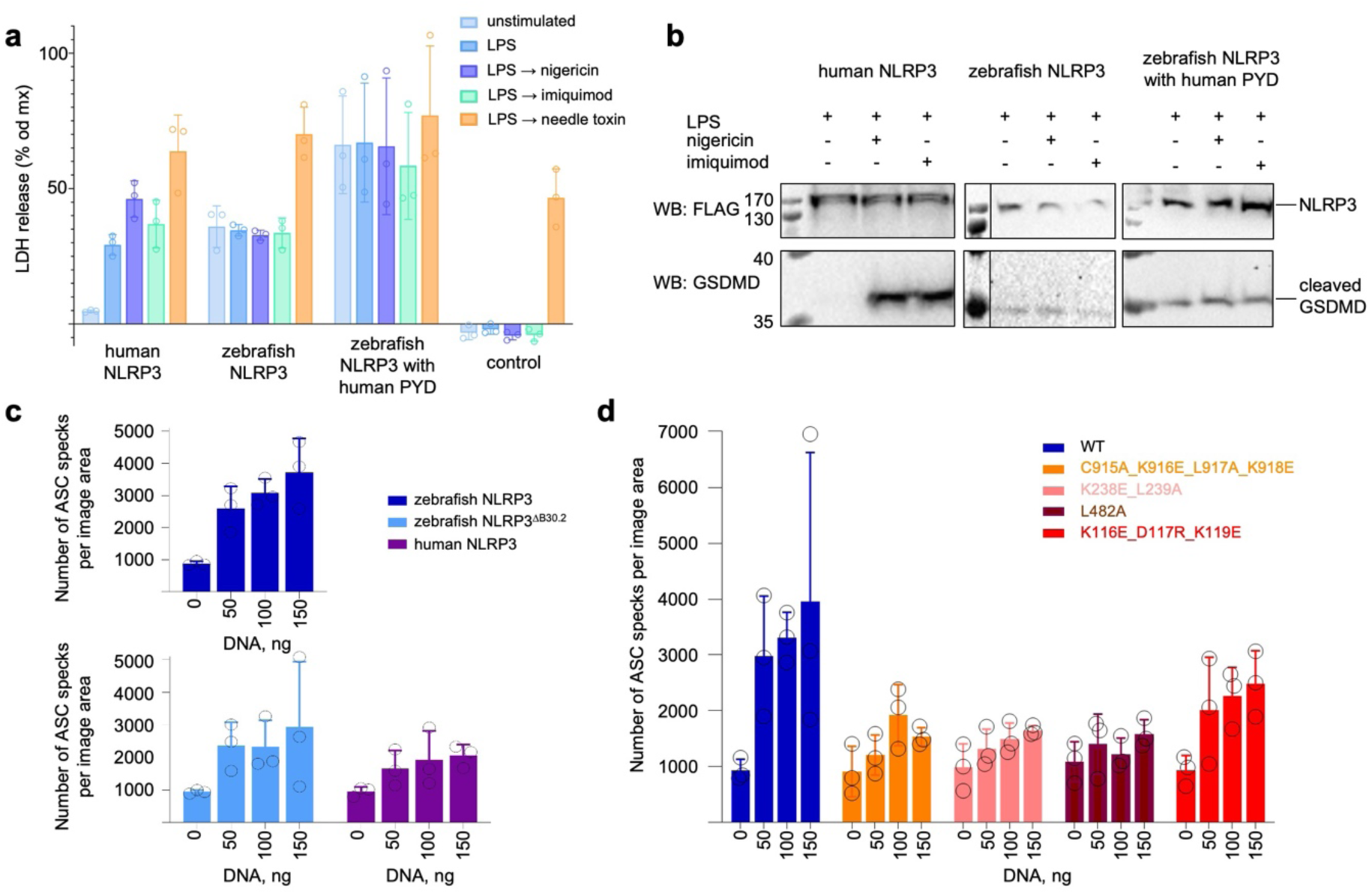
Zebrafish NLRP3 forms an active inflammasome in cells. **a**, LDH release assay in NLRP3-KO BlaER1 cells reconstituted with the indicated FLAG-tagged NLRP3 constructs or control plasmid. Expression was induced with doxycycline (Dox) for 18 h. Cells were primed with 200 ng/ml LPS for 4h followed by activation with 6.5 µM nigericin, 30 µg/ml imiquimod or 0.025 mg/ml needle toxin for 2 h. Data are represented as mean ± SD, n = 3. **b**, Western blots of the whole cell lysates from THP-1 NLRP3-KO cells reconstituted with human or zebrafish NLRP3 or zebrafish NLRP3 containing human PYD. Cells were treated with 1 µg/ml LPS for 3 h followed by activation with 20 µM nigericin or 200 µM imiquimod for 1 h. FLAG-tagged NLRP3 constructs and cleaved GSDMD were visualized with corresponding antibodies. **c, d,** ASC speck formation in HEK293T cells stably expressing YFP-ASC, upon transfection with indicated NLRP3 constructs: human, zebrafish NLRP3 and zebrafish NLRP3 with B30.2-domain deletion (zebrafish NLRP3^ΔB30.2^) (**c**); wild-type and oligomer-disrupting mutants of zebrafish NLRP3 (**d**). Cells were transfected with the indicated amount of DNA, and the number of ASC specks per image area was calculated 24 h post-transfection. Data are represented as mean ± SD, n = 3.

Such spontaneous activation at high expression levels has also been observed for the human NLRP3 inflammasome expressed in HEK293T cells^60^. Therefore, we decided to utilize overexpression in HEK293T to compare various NLRP3 constructs. For that purpose, we generated a HEK293T cell line stably expressing YFP-tagged human ASC at a relatively low level to avoid NLRP3-independent ASC speck formation and transfected it with increasing amounts of NLRP3 constructs. Inflammasome formation was assessed by the formation of ASC specks – a well-established readout for inflammasome activation^61,62^. Like human NLRP3, both wild-type and B30.2 domain-deleted (NLRP3^ΔB30.2^) zebrafish NLRP3 induced ASC speck formation upon overexpression, in agreement with previous reports^36^ (Fig. 5c, Supplementary Fig. 9b). To address whether ASC speck formation was triggered by the active NLRP3 complex and not by unspecific protein aggregation, we repeated this experiment using zebrafish NLRP3 mutants disrupting the active 21-mer oligomer (Fig. 2d). Indeed, disruption of any of the four interfaces involved in active 21-mer zebrafish NLRP3 formation (Fig. 2a, e, f) reduced zebrafish NLRP3 activity in triggering ASC speck formation upon overexpression in the YFP-ASC expressing HEK293T cell line, even when expression levels of some mutants were higher than those of the wild-type protein (Fig. 5d, Supplementary Fig. 9c). Taken together, our data demonstrate that zebrafish NLRP3 can form an active inflammasome in human cells and requires its unique 21-mer conformation for signaling.

### Zebrafish NLRP3 does not form inactive oligomeric “cages” and does not co-localize with the Golgi

One of the very important steps in a canonical activation of human NLRP3 is the formation of the membrane-bound NLRP3 “cages”^11–13^. Despite collecting zebrafish NLRP3 from the heavy sucrose gradient fractions (Fig. 1b), cryo-EM data processing did not reveal any additional NLRP3 species, which could resemble a zebrafish NLRP3 “cage” (Fig. 1d, Supplementary Fig. 1c). Given that zebrafish NLRP3 has a very short PYD-FISNA linker (Fig. 3d, Supplementary Fig. 6e), we hypothesized that it might prevent PYD positioning needed for “cage” stabilization. However, an insertion of the human linker into zebrafish NLRP3 (Supplementary Fig. 10a) neither lead to “cage”, nor to an active 21-mer formation (Fig. 6a, Supplementary Fig. 10b). Sequence alignment furthermore revealed that both “face-to-face” and “back-to-back” interfaces responsible for human or mouse NLRP3 “cage” formation are not conserved in zebrafish NLRP3 (Supplementary Fig. 10c) and would lead to steric clashes between zebrafish NLRP3 monomers (Fig. 6b, c).

**Fig. 6.**
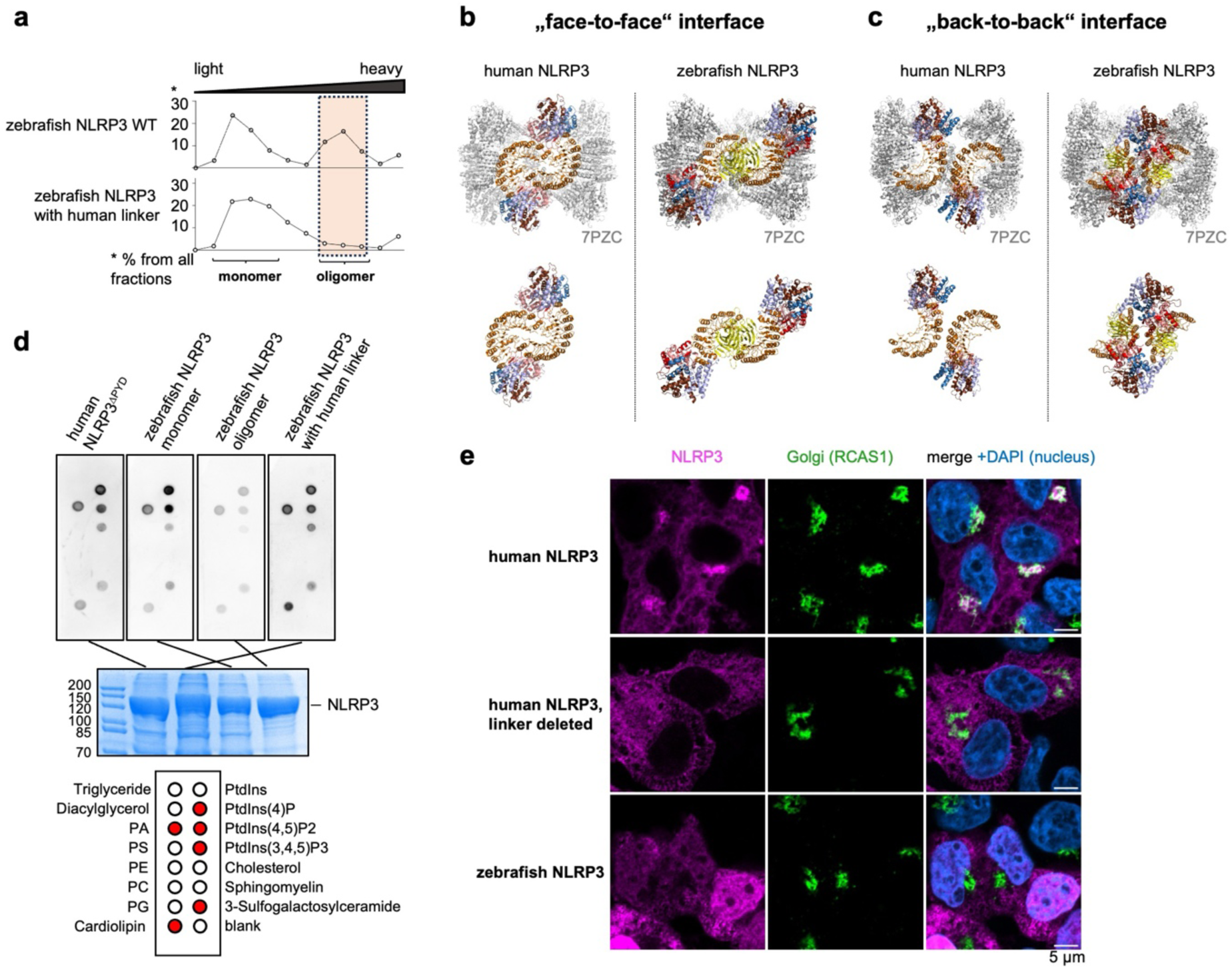
Zebrafish NLRP3 activity is “cage”- and TGN-independent. **a**, Sucrose gradient profiles of zebrafish NLRP3 WT and zebrafish NLRP3 containing the human PYD-FISNA linker sequence. The profiles were calculated by quantification of zebrafish NLRP3 bands from the SDS-PAGE images in Supplementary Fig. 10b. The fractions containing NLRP3 oligomer are highlighted in orange. **b, c**, Superposition of inactive zebrafish NLRP3 (colored by domains) with “face-to-face” (**b**) and “back-to-back” (**c**) interfaces forming human NLRP3 “cage” (grey, PDB: 7PZC). **d**, *In vitro* lipid blot assay of human monomer (ΔPYD), zebrafish monomer and active oligomer NLRP3, and zebrafish NLRP3 containing the human linker sequence. In the middle, SDS-PAGE with loading controls. On the bottom, lipid scheme. Lipids interacting with NLRP3 constructs are highlighted in red. **e**, Confocal microscopy imaging of HEK293T cells reconstituted with mScarlet-tagged human WT, human linker-deleted, and zebrafish WT NLRP3 constructs for NLRP3 (mScarlet, magenta), Golgi protein RCAS1 (IF, green), and DNA (Hoechst dye, blue). Scale bar 5 µm.

Since zebrafish NLRP3 did not form “cages”, we wondered whether it could still bind membranes and co-localize with the Golgi, as human NLRP3 does^11,23,31,63^. As reported before, the exon 3-encoded linker region of human NLRP3 is critical for both lipid binding and association with the Golgi^28,63^, and its deletion renders human NLRP3 cytosolic^28^. Since this linker region is missing in zebrafish NLRP3 (Supplementary Fig. 6e, Supplementary Fig. 10a), it is tempting to hypothesize that zebrafish NLRP3 might demonstrate similar behavior. Unexpectedly, *in vitro* lipid blot assays, however, showed that both monomeric and active oligomeric zebrafish NLRP3 bound to the same mono-, di- and tri-phosphorylated phosphatidylinositides and cardiolipin, like human NLRP3, though the active 21-mer showed reduced lipid binding in comparison to a monomer (Fig. 6d). Interestingly, an insertion of the human linker region, which harbors critical palmitoylation sites^30,31^, into zebrafish NLRP3 (Supplementary Fig. 10a) did not increase its lipid binding *in vitro* (Fig. 6d). This likely emerges from significant differences between human and zebrafish NLRP3, which would prohibit human palmitoylation enzymes to efficiently modify the linker in the chimeric construct.

We next aimed to address the localization of zebrafish NLRP3 in cells. For this purpose, HEK293T cell lines stably expressing N-terminally mScarlet-tagged human and zebrafish NLRP3 constructs were generated. Remarkably, despite demonstrating the same lipid binding *in vitro* as human NLRP3, zebrafish NLRP3 did not co-localize with the Golgi based on confocal microscopy images (Fig. 6e). Such a dramatic difference in localization between human and zebrafish NLRP3 might, at least partially, emerge from differences in their palmitoylation sites. For human NLRP3, palmitoylation of C130 and C958 was shown to be critical for its Golgi localization^30,31^. These residues, however, are not conserved in zebrafish NLRP3, which might explain the lack of Golgi localization (Supplementary Fig. 10d). Instead, zebrafish NLRP3 was recently shown to undergo a C1037 palmitoylation in the B30.2 domain, which was critical for its activation^35^. Thus, while both zebrafish and human NLRP3 likely rely on palmitoylation and membrane binding, the source of the membranes involved in their regulation may differ between species.

### Zebrafish NLRP3 does not interact with NEK7 and forms MTOC-independent ASC specks

Since zebrafish NLRP3 neither forms “cages” nor co-localizes with the Golgi, we were wondering whether it still associates with NEK7. The LRR interface involved in NEK7 binding is occupied by a C-terminal B30.2 domain in zebrafish NLRP3, therefore, an additional binding to NEK7 would result in a steric clash and thus is very unlikely (Fig. 7a). Sequence analysis revealed that the NEK7-interacting residues of human NLRP3 are not conserved in zebrafish (Supplementary Fig. 11). To address NEK7 binding experimentally, we co-expressed FLAG-tagged NLRP3 constructs with HA-tagged human or zebrafish NEK7 in Expi293F cells and analyzed the binding using FLAG-beads pull-down. Indeed, zebrafish NLRP3 did not interact with human or zebrafish NEK7, while human NLRP3 and its exon 3-deleted version could bind both NEK7 constructs (Fig. 7b). Interestingly, a deletion of the B30.2 domain enabled zebrafish NEK7 binding by zebrafish NLRP3 (Fig. 7b), despite significant differences in LRR sequence compared to human NLRP3. Thus, although zebrafish NLRP3 does not engage with NEK7 due to a hindrance from the B30.2 domain, its LRR interface itself is compatible with NEK7 binding.

**Fig. 7.**
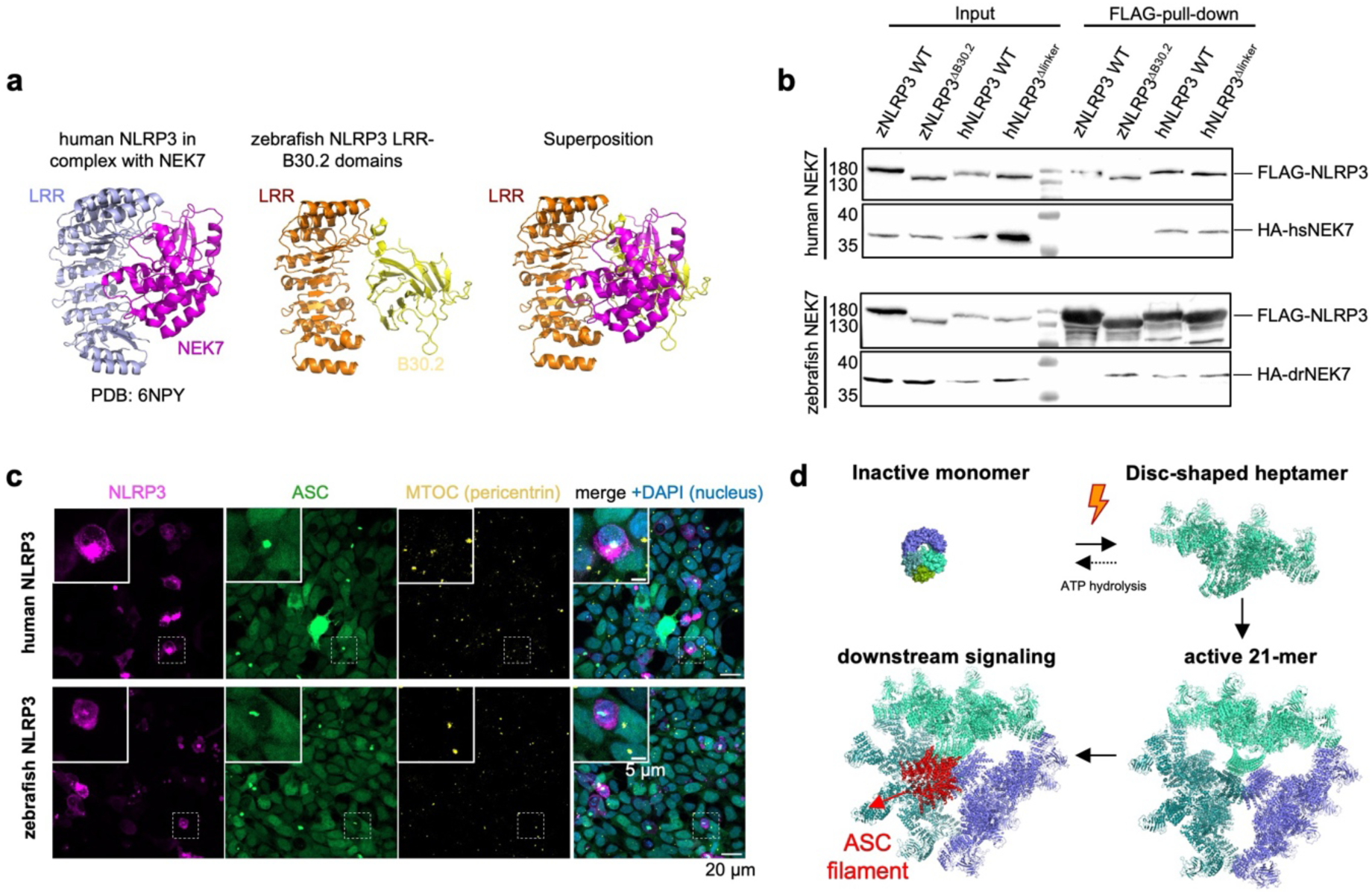
Zebrafish NLRP3 undergoes a NEK7 and MTOC-independent activation. **a**, A comparison of the NEK7-bound LRR domain of human NLRP3 (blue, PDB: 6NPY) with LRR-B30.2 domains of zebrafish NLRP3 (orange and yellow, respectively). NEK7 is sown in magenta. **b**, The FLAG pull-down from Expi293F cells transfected with the HA-tagged human or zebrafish NEK7 and FLAG-tagged NLRP3 constructs analyzed by Western blot using anti-FLAG and anti-HA antibodies. **c**, Confocal microscopy imaging of HEK293T cells reconstituted with EGFP-tagged ASC and transfected with mScarlet-tagged zebrafish or human WT NLRP3 for NLRP3 (mScarlet, magenta), ASC (EGFP, green), MTOC marker pericentrin (IF, yellow), and DNA (Hoechst dye, blue). Scale bar 20 µm and 5 µm for the field view and the close-up, respectively. **d**, Proposed zebrafish NLRP3 activation model. Upon activation, an inactive monomeric NLRP3 converts to a single disc-shaped heptamer, which undergoes a second oligomerization step to form an active 21-mer oligomer capable of ASC oligomerization.

As zebrafish NLRP3 does not bind NEK7, we were wondering whether it forms an active inflammasome independently of MTOC localization, like observed for linker-deleted human NLRP3^28^, NLRC4 and AIM2^14^. For this purpose, mScarlet-tagged human and zebrafish NLRP3 constructs were transfected into HEK293T cells stably expressing human EGFP-tagged ASC, and their localization was observed using confocal microscopy (Fig. 7c). Interestingly, under the tested conditions both zebrafish and human NLRP3 induced ASC speck formation distally from the MTOC (Fig. 7c).

Collectively, our data demonstrate that zebrafish NLRP3 forms an active inflammasome in human cells yet lacks the major hallmarks of the canonical NLRP3 activation pathway. Thus, zebrafish NLRP3 engages in a “cage- “, Golgi-, NEK7- and MTOC-independent non-classical activation pathway, which would explain the unusual architecture of its active oligomeric form, and offers an alternative structural mechanism of NLRP3 activation.

## Discussion

Our discovery of a novel mode of NLRP3 activation utilized by zebrafish NLRP3 offers a previously unknown mechanism of NLRP3 activation (Fig. 7d). According to our hypothesis, activation of this ancient NLRP3 homolog involves two steps of oligomerization, like for human NLRP3, but the oligomers formed are drastically different: At the first step, instead of inactive membrane-bound oligomeric “cages”^11–13^ formed by human NLRP3, zebrafish NLRP3 directly forms disc-shaped oligomers (Fig. 2c). These oligomers are stabilized by NACHT domains (Fig. 2e, f), which are present in an “open” ATP-bound state (Fig. 4b, Supplementary Fig. 3c, Supplementary Fig. 7a) as in the active NLRP3 inflammasome disc^15^. These zebrafish NLRP3 disc-shaped oligomers alone, however, cannot nucleate ASC oligomerization and trigger downstream signaling. Though 7 PYDs should be enough to form one helical turn (6 PYDs) of the PYD filament, their reduced flexibility prohibits filament formation. Indeed, zebrafish NLRP3 PYDs are connected to the disc-shaped FISNA-NACHT-LRR-B30.2 heptamers through very short 9-aa linkers as opposed to 41-aa-long flexible linkers in human NLRP3 (Fig. 3d, Supplementary Fig. 6d, e). Thus, zebrafish heptamers must undergo a second round of oligomerization into a trimer of heptamers to form a short functional PYD segment capable of downstream signaling (Fig. 3d, e). Of note, the human apoptosome receptor Apaf-1 also forms an active heptamer but does not require further oligomerization due to sufficiently long linkers connecting the CARD domains to the rest of the structure^64,65^. Additional oligomerization of oligomeric discs is thus a completely novel mode of inflammasome activation, which, according to our knowledge, has not been reported before. It broadens our knowledge of NLRP3 plasticity, which can adopt various forms – from hexamers^13,66^ and octamers^66^ to decamers and dodecamers “cages”^11–13^ and 10-/11-fold discs^15^ – and expands the wide range of unique inflammasome activation mechanisms discovered so far^67^.

Despite being distant homologs of human NLRP3, various teleost NLRP3, including zebrafish NLRP3^36,37^, have been shown to respond to canonical NLRP3 activation stimuli^33–35^. However, under the conditions tested in this study, zebrafish NLRP3 formed active inflammasomes in BlaER1, THP-1, and HEK293T cells spontaneously, and a strictly stimulus-dependent activation could not be observed (Fig. 5a-c). This explains how the active 21-mer zebrafish NLRP3 could be obtained from human cells in the absence of NLRP3 activators, which were required to obtain active human NLRP3 disc^15^. However, since human cell lines were tested, numerous factors including potential incompatibility of co-factors and modification enzymes and expression temperature, which is higher for human in comparison to zebrafish cell cultures, could have contributed to zebrafish NLRP3 hyperactivity.

Importantly, the ability of zebrafish NLRP3 to form an inflammasome must emerge from active 21-mer zebrafish NLRP3 oligomers rather than non-specific aggregation. Indeed, no significant aggregation was observed in HEK293T cells using confocal microscopy (Fig. 6e) and mutations disrupting the 21-mer arrangement abolished zebrafish NLRP3 activity in HEK293T cells (Fig. 5d). One cannot assume, however, that missing zebrafish factors alone are responsible for zebrafish NLRP3 activation. Indeed, apart from the active 21-mer oligomeric zebrafish NLRP3 form, a significant amount of monomer was present during purification (Fig. 1b), suggesting that some activating monomer-to-oligomer transition must happen in human cell lines. Since immortalized cell lines in general have impaired checkpoints and metabolic distortions in comparison to primary cells, these homeostasis disturbances might have been sufficient for the zebrafish NLRP3 activation^2^. Thus, zebrafish NLRP3 might be a promising model to search for the precise NLRP3-activating event, which remains elusive.

Like for human NLRP3, ATP binding plays an important role in stabilizing the active zebrafish NLRP3 21-mer conformation (Fig. 4a-c). Consequently, mutation in a “Glu-switch” region, which is critical for ATP hydrolysis, drastically increased the amount of active zebrafish NLRP3 21-mer while retaining the same structural arrangement (Fig. 4f-j). We do not know whether other disease mutations would have the same effect, calling for further experimental elucidation. Interestingly, binding of an unhydrolyzable ATP homolog to a zebrafish NLRP3 monomer was sufficient to induce its multimerization (Fig. 4c) but not in the form of the specific active complex. This might indicate that a general constraint for multimerization is relieved by nucleotide exchange, but that other cellular constraints, like scaffolding membranes^11,23,68^, are additionally needed for the assembly of the highly ordered active zebrafish NLRP3 complex.

Surprisingly, while zebrafish NLRP3 bound the same lipids as human NLRP3 *in vitro* (Fig. 6d), it did not co-localize with the Golgi (Fig. 6e). Palmitoylation and membrane binding were recently shown to be critical for teleost NLRP3 activation, though a different palmitoylation site in the B30.2 domain was proposed^35^. Due to significant differences in palmitoylation sites in comparison to human NLRP3 (Supplementary Fig. 10d) it is thus likely that zebrafish and human NLRP3 display different preferences for membranes or lipids. Indeed, clustering on various membranes can potentiate human NLRP3 activation, as shown recently^68^. Interestingly, while human NLRP3 has been proposed to interact with membranes predominantly in the “cage” form, it cannot be excluded that other NLRP3 conformations, including monomers, can also interact with lipids^11,12^. Indeed, both zebrafish and human^11^ NLRP3 monomers can bind lipids (Fig. 6d). Remarkably, the zebrafish NLRP3 monomer demonstrated a much stronger interaction with lipids than a 21-mer active oligomer (Fig. 6d). Such loss of lipid interaction upon activation has also been proposed for human NLRP3 upon its phosphorylation with Bruton’s tyrosine kinase, which was shown to increase NLRP3 activity^69^. Thus, a transient membrane interaction – in the form of a monomer or an oligomer – may be a common consensus in NLRP3 activation across the evolution.

With human NLRP3-specific hallmarks like “cage” formation (Fig. 6b, c) and Golgi localization (Fig. 6e) missing in zebrafish NLRP3, it is not surprising that zebrafish NLRP3 also does not interact with NEK7 (Fig. 7b) or depend on MTOC, where NEK7 typically resides^25–27^, for ASC speck formation (Fig. 7c). This suggests that some non-classical activation path must be taking place. The zebrafish NLRP3 activation pathway shares numerous similarities with a recently described Golgi- and MTOC-independent human NLRP3 pathway, discovered by deletion of the flexible PYD-FISNA linker encoded by human *NLRP3* exon 3, which still forms a functional inflammasome^28^. Indeed, like zebrafish NLRP3, human exon 3-deleted NLRP3 did not form “cages”, did not co-localize with the Golgi (also in Fig. 6e), and did not require the MTOC for active inflammasome formation^11^. Mechanistically, in this construct, neither a canonical 10-/11-fold disc^15^ nor a heptamer would suffice for PYD filament formation due to limited flexibility in PYD positioning, similar to that assumed for zebrafish NLRP3. Therefore, an additional step of NLRP3 discs oligomerization may be required for non-“cage” human NLRP3 for functional inflammasome assembly, as observed for zebrafish NLRP3. We do not know whether a zebrafish-like activation pathway takes place in human cells under physiological conditions. However, its compatibility with human inflammasome components and similarity to the human MTOC-independent NLRP3 pathway are striking. The ability of zebrafish NLRP3 LRR to bind NEK7 suggests NEK7 binding (Fig. 7b), as well as the PYD-FISNA linker acquired in vertebrates, to be more important for NLRP3 regulation than for activation itself.

Collectively, using a structural biology and biochemistry approach, we show a new non-canonical mechanism of NLRP3 oligomerization and activation utilized by zebrafish NLRP3 and propose zebrafish NLRP3 as a promising model for searching for key NLRP3 activating events and alternative activation strategies.

---

**Supplementary Fig. 1.**
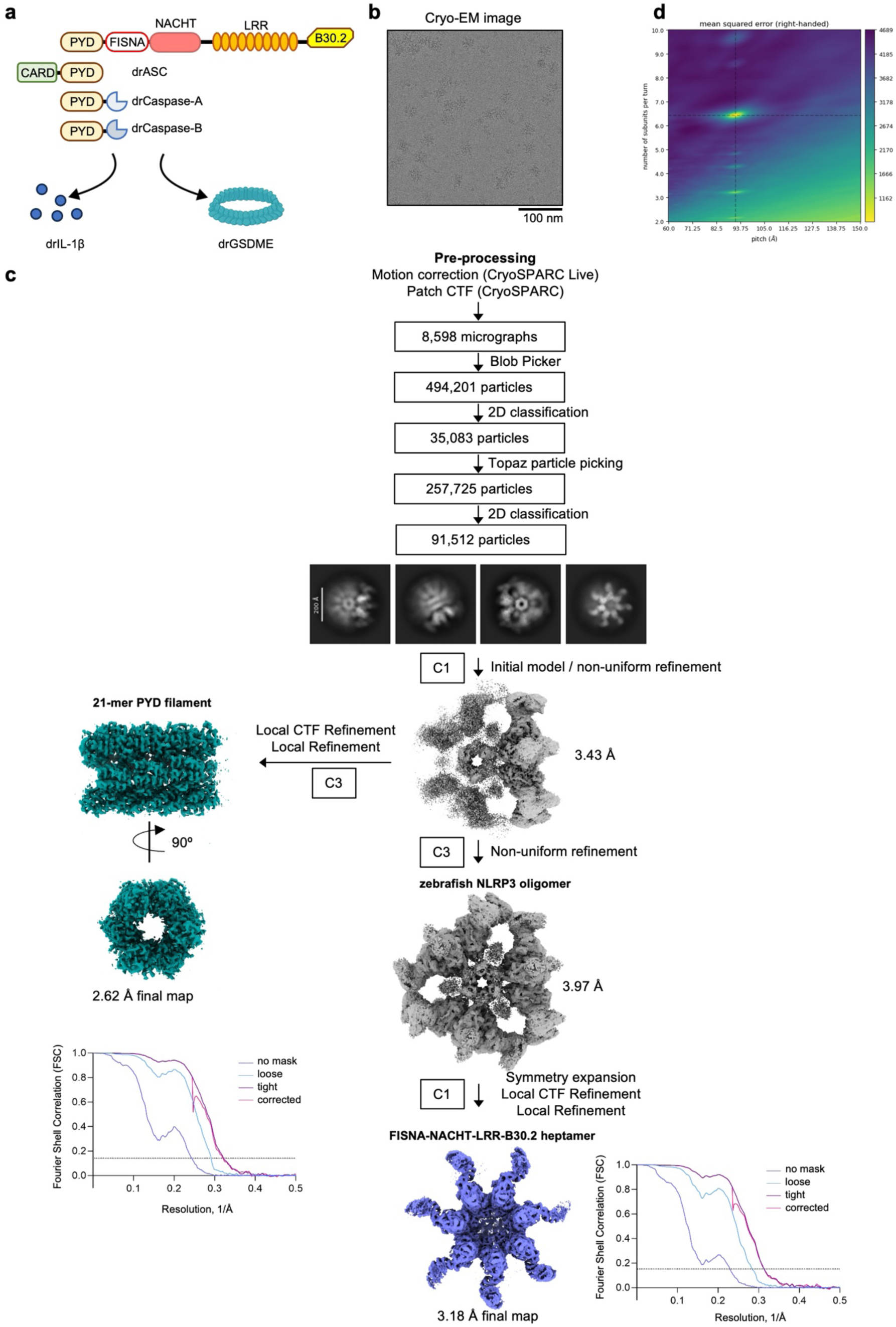
Structure determination workflow for the WT zebrafish NLRP3 oligomer. **a**, Schematic representation of the zebrafish NLRP3 inflammasome complex and its activation outcomes. Inflammasome components are depicted with their domain structure. **b**, A representative cryo-EM micrograph. **c**, Structure determination workflow with the cryo-EM map for the full zebrafish NLRP3 complex and final cryo-EM maps of a FISNA-NACHT-LRR-B30.2 domains-containing heptamer and PYD 21-mer, and corresponding Fourier Shell Correlation (FSC) curves. The dashed line indicates the 0.143 cutoff. **d**, 2D plot of the mean squared error surface along the helical pitch and number of subunits per turn. The best estimate of helical symmetry parameters is located on the interception of the global minima depicted with dashed lines. See also Supplementary Table 1.

**Supplementary Fig. 2.**
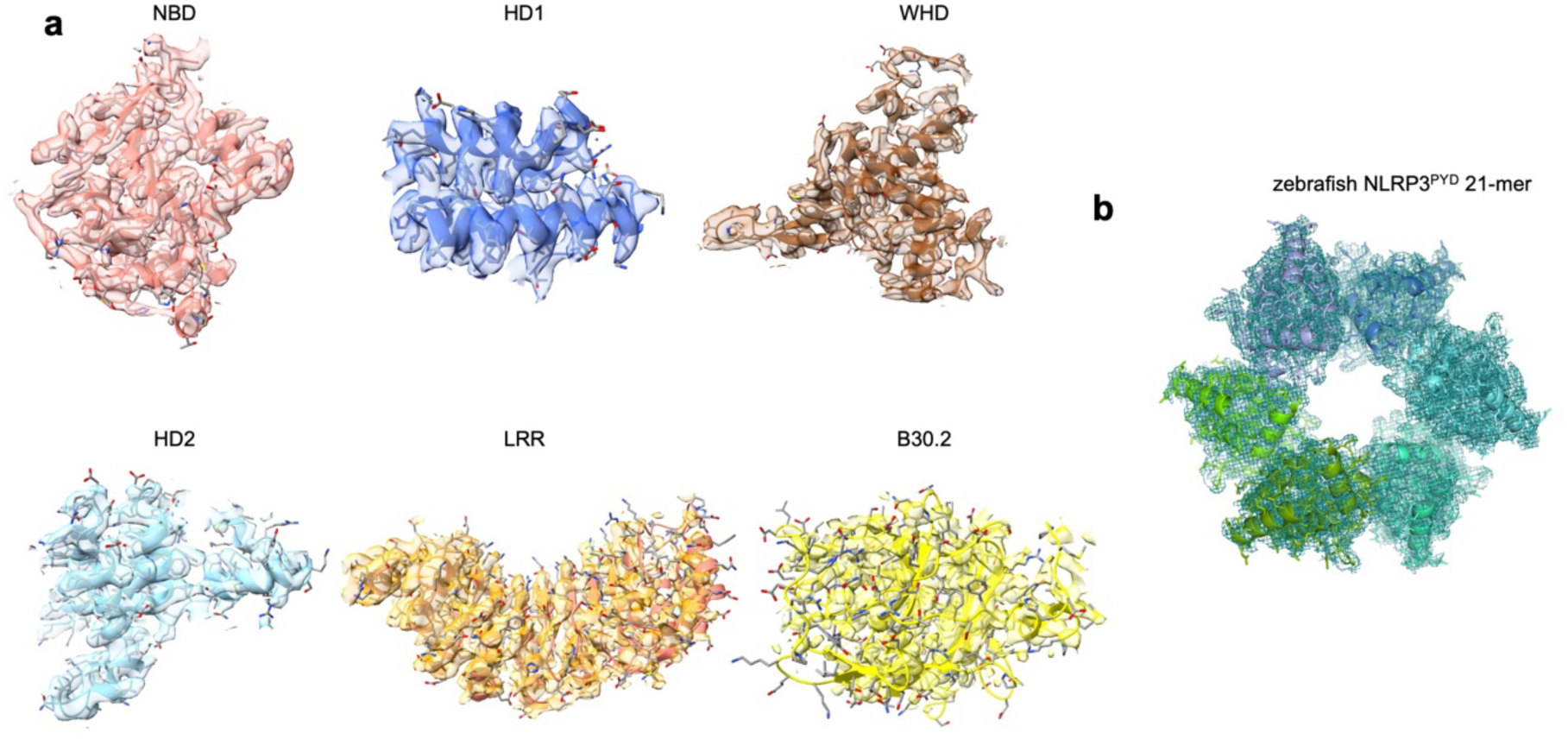
Cryo-EM map quality of zebrafish NLRP3 regions. **a, b**, Cryo-EM maps of various subdomains of the WT zebrafish NLRP3 oligomer shown as isomesh at 1 root mean square deviation (RMSD) with a 2.5 Å radius.

**Supplementary Fig. 3.**
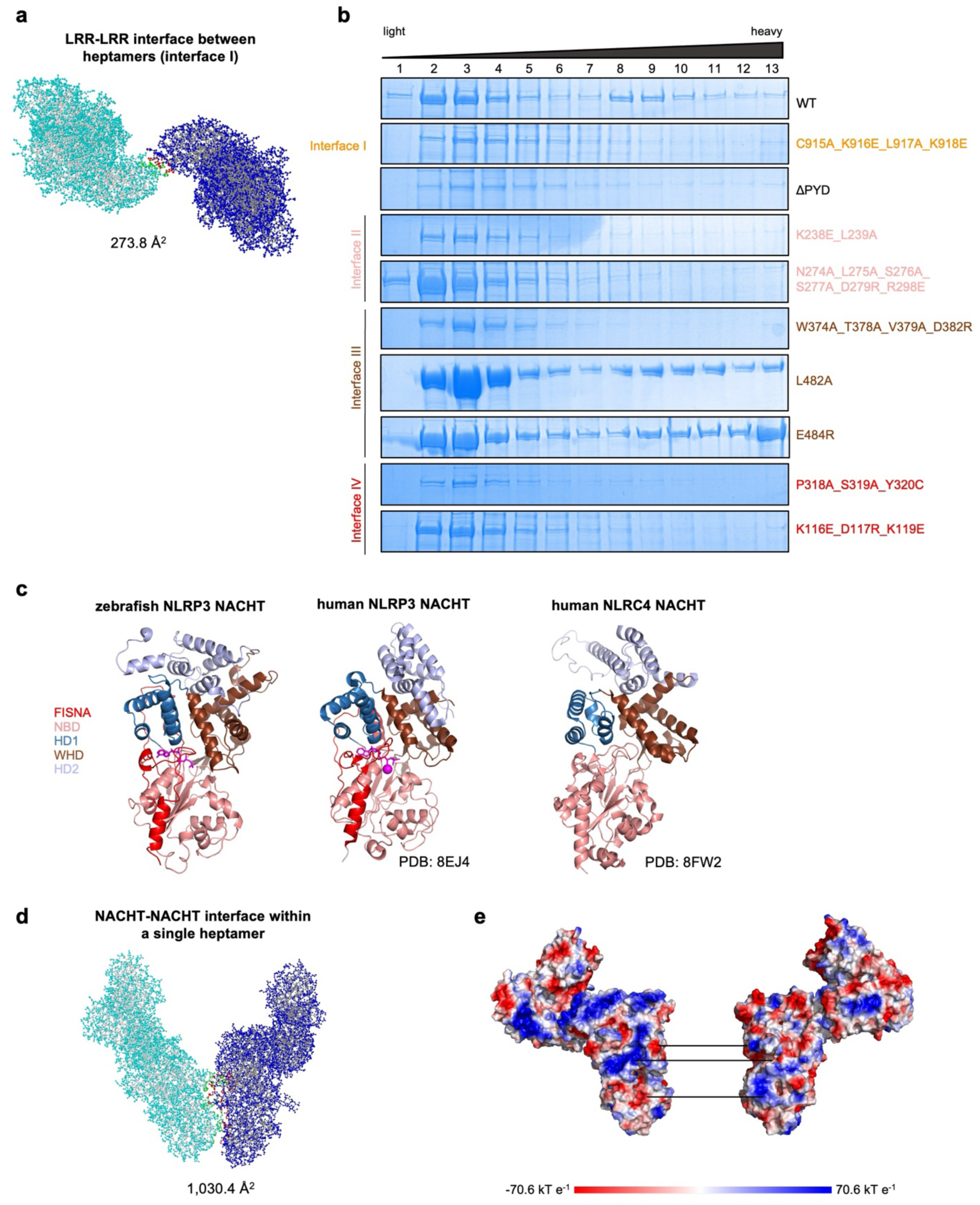
Analysis of the main interfaces within the WT zebrafish NLRP3 oligomer. **a**, Ball-and-stick model of the zebrafish NLRP3 monomers (blue and cyan) from two different heptamers interacting with their LRR domains in trans. Interface regions are defined by PDBePISA^46^ and colored red and green. **b**, SDS-PAGE gels of sucrose gradient fractions of zebrafish NLRP3 WT and interface mutants, color-coded by interface: orange for interface I, salmon for interface II, brown for interface III, and red for interface IV. **c**, Comparison of zebrafish NLRP3 NACHT with NACHT domains of the active NLRP3 (PDB: 8EJ4) and NLRC4 (PDB: 8FW2) disc structures. **d**, Ball-and-stick model of the adjacent monomers (blue and cyan) within the zebrafish NLRP3 oligomer. Interface regions are defined by PDBePISA^46^ and colored red and green. **e**, Surface electrostatic potential of the two zebrafish NLRP3 monomers, shown in the range of red (-70.6 kT/e, negatively charged) to blue (70.6 kT/e, positively charged). Surface charge complementarity between the interacting monomers is indicated with lines.

**Supplementary Fig. 4.**
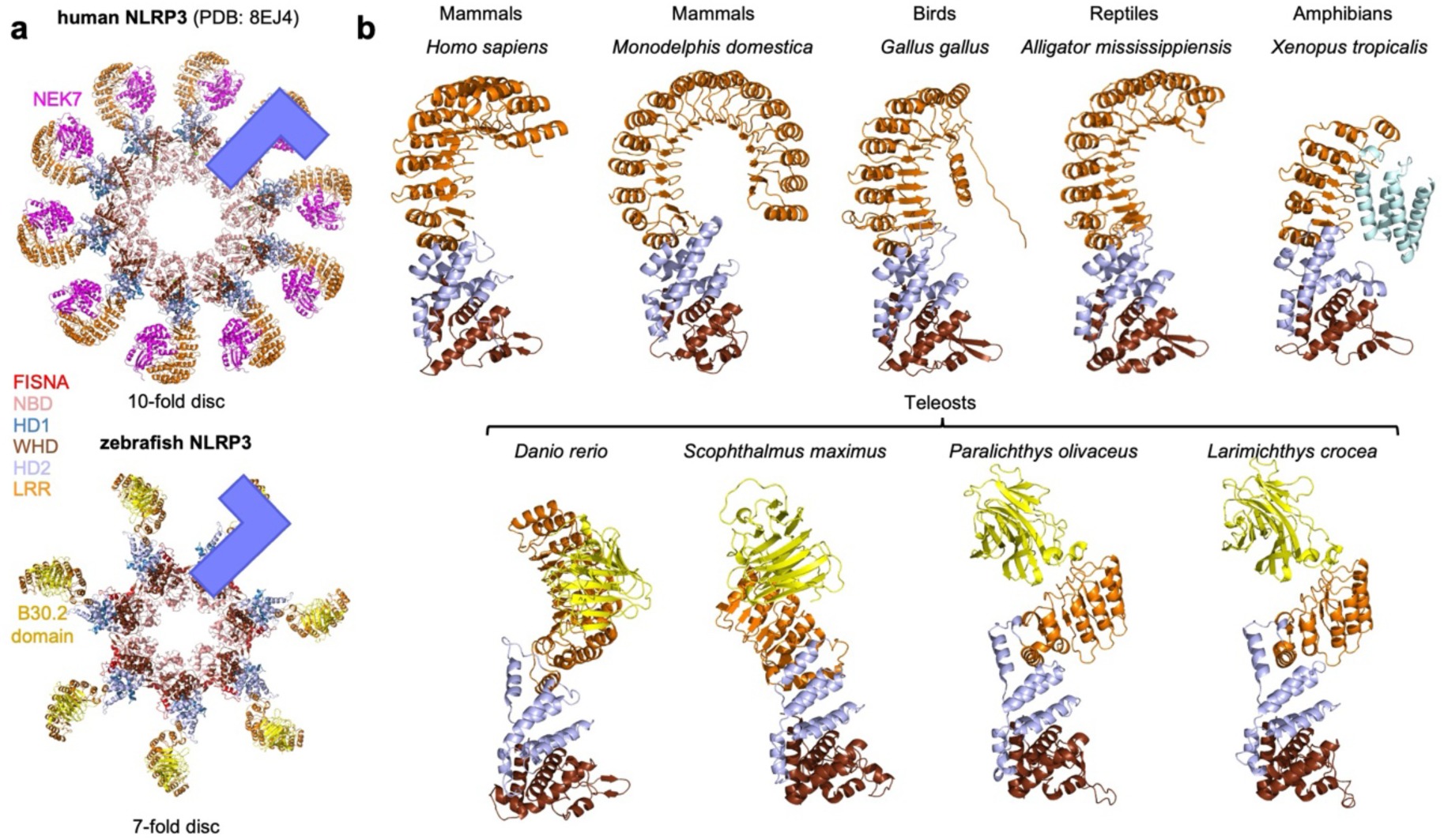
Orientations of the LRR-B30.2 domains in vertebrata. **a**, Atomic models of human and zebrafish FISNA-NACHT-LRR oligomeric discs, colored by domains. Orientation of the LRR relatively to the NACHT is indicated with a blue L-shaped figure. **b**, AlphaFold 3^70^ prediction models of NLRP3 from various vertebrate species aligned on WHD-HD2 segment: *M. domestica* XP_007491161.2, *G. gallus* NP_001335876.2, *A. mississippiensis* XP_019340765.1, *X. tropicalis* XP_017951762.1, *D. rerio* MN088121.1, *S. maximus* AWO98340.1, *P. olivaceus* XP_069375685.1, *L. crocea* XP_010744826.3, and a structure of human NLRP3 - PDB: 8EJ4. WHD-HD2-LRR and WHD-HD2-LRR-B30.2 domains are shown, colored by domain.

**Supplementary Fig. 5.**
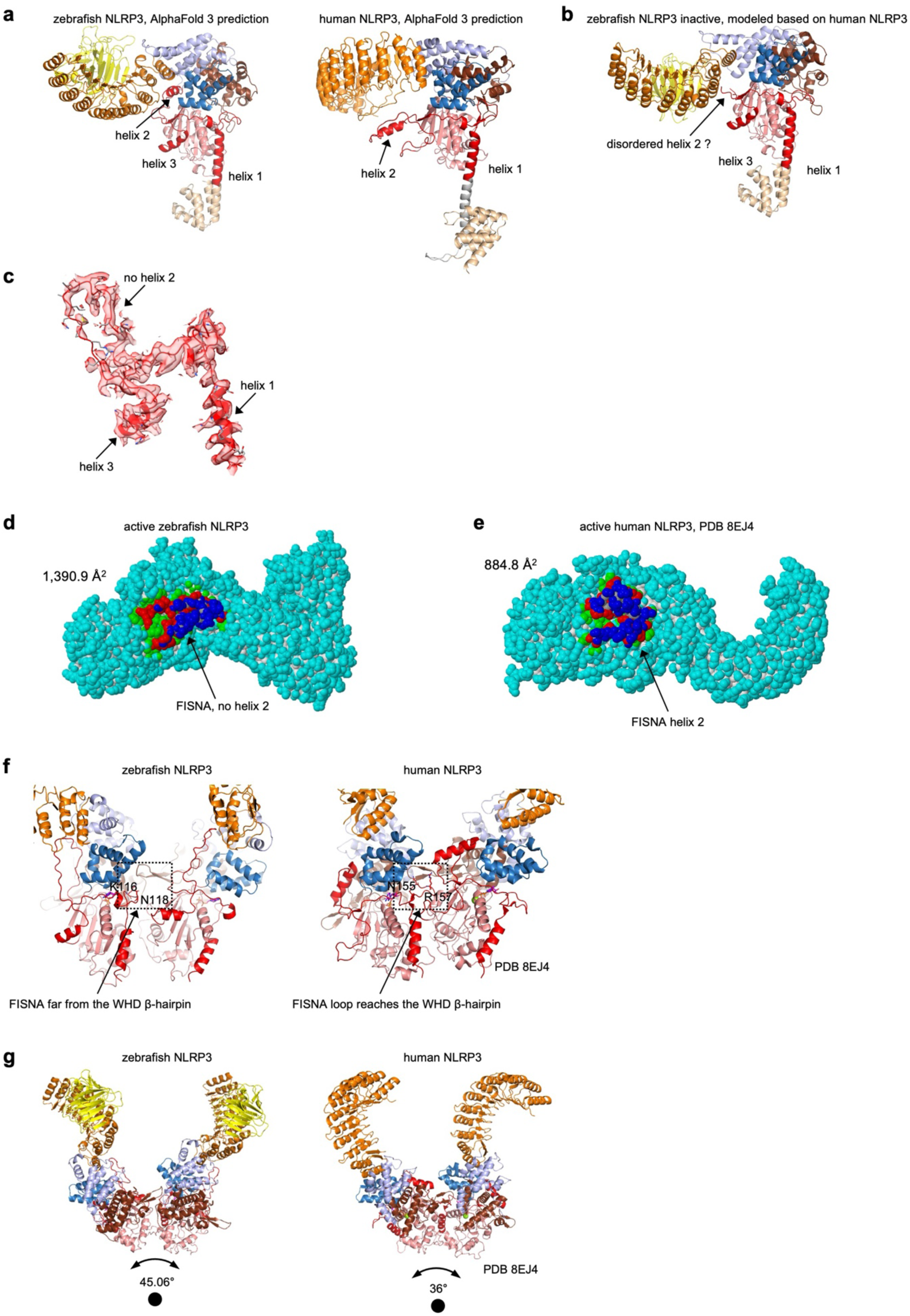
Conformation of the FISNA domain in the WT zebrafish NLRP3 oligomer. **a**, AlphaFold 3^70^ prediction models of zebrafish (left) and human (right) NLRP3, colored by domain. FISNA α-helices are numbered and indicated in red. **b**, Inactive zebrafish NLRP3 modeled based on inactive human NLRP3 (PDB: 7PZC), colored by domain. **c**, Cryo-EM map of FISNA domain of the active zebrafish NLRP3 monomer, shown as isomesh at 1 root mean square deviation (RMSD) with a 2.5 Å radius. **d, e**, Ball-and-stick model of the zebrafish (**d**) and human (**e**) NLRP3 monomer in active conformation. “Helix 2” FISNA region is blue, the rest of the NLRP3 molecule is cyan. Interface regions are defined by PDBePISA^46^ and colored red and green. **f**, An overview of adjacent FISNA-NACHT domains of zebrafish (left) and human (right) NLRP3 in the active conformation, colored by domains. Residues N155 and R157 involved in NACHT-NACHT interactions in the human NLRP3 oligomeric disc^15^ and corresponding residues in zebrafish NLRP3 are shown as sticks. **g**, Atomic models of two adjacent NLRP3 monomers from active zebrafish (left) and human (right) oligomeric discs, colored by domains. The rotation degree between two monomers is indicated.

**Supplementary Fig. 6.**
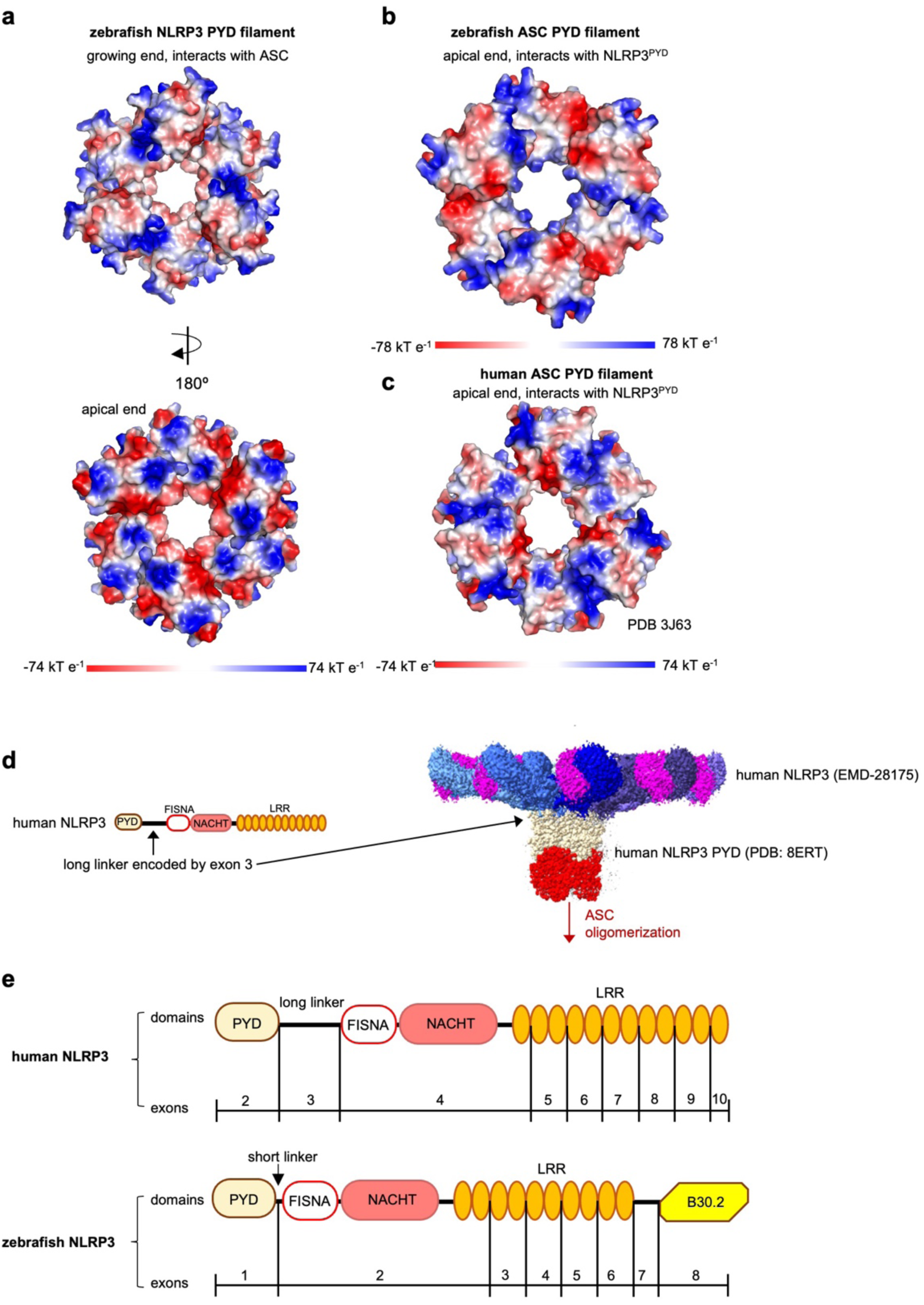
Position and surface charge of zebrafish and human NLRP3 PYD filament within the corresponding active NLRP3 structures. **a-c**, Top views of the 21-mer zebrafish NLRP3 (**a**), zebrafish ASC (AlphaFold 3^70^ prediction) (**b**) and human ASC (PDB: 3J63) (**c**) PYD filaments colored by surface electrostatic potential shown in range of red (negatively charged) to blue (positively charged). The apical ends of ASC^PYD^ filaments interacting with NLRP3^PYD^ are shown. **d** Domain structure of human NLRP3 with long PYD-FISNA linker encoded by the exon 3 (left); cryo-EM map of the active NLRP3 disc colored by monomer (EMDB-28175) and atomic model of the human NLRP3 PYD filament (PDB: 8ERT, right). PYD domains of active NLRP3 are colored in wheat. NLRP3 PYD domains, which shall correspond to ASC in the inflammasome assembly, are colored red. The proposed direction of ASC oligomerization is indicated with a red arrow. **e**, Exon structure and corresponding domain architecture of human and zebrafish NLRP3.

**Supplementary Fig. 7.**
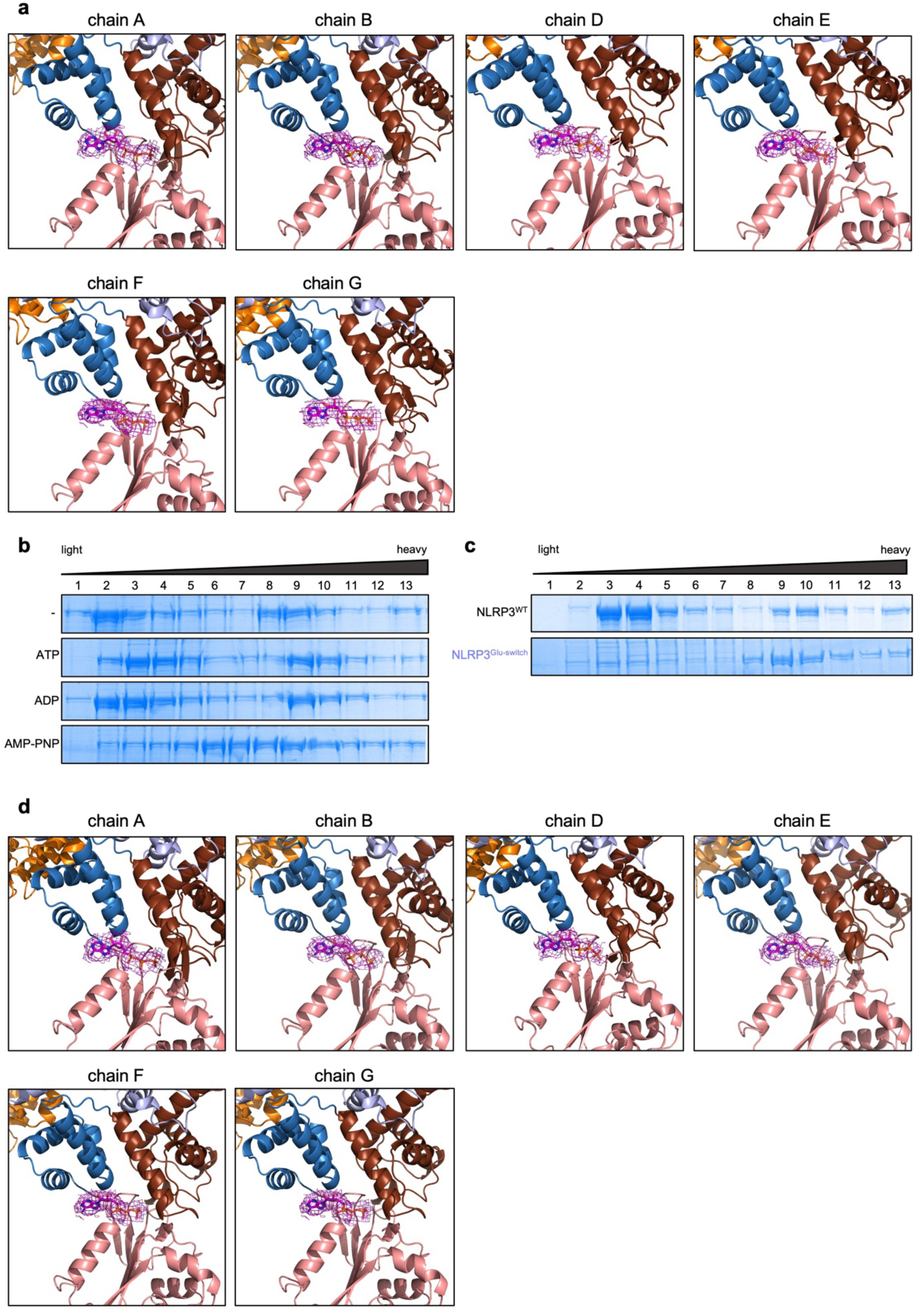
Binding of ATP and its analogs by zebrafish NLRP3. **a**, A detailed view of the ATP-binding pockets in all protein chains within the wild-type zebrafish NLRP3 heptamer. Cryo-EM density of ATP is shown as isomesh at 1 root mean square deviation (RMSD) with a 2.5 Å radius, in magenta. The respective representation of chain C is shown in Fig. 4b. **b**, SDS-PAGE gels of sucrose gradient fractions of zebrafish NLRP3 WT purified without or in the presence of the indicated nucleotides. **c**, SDS-PAGE gels of sucrose gradient fractions of WT and “Glu-switch”-mutated zebrafish NLRP3. **d**, A detailed view of the ATP-binding pockets in all protein chains within the “Glu-switch”-mutated zebrafish NLRP3 heptamer. Cryo-EM density of ATP is shown as isomesh at 1 root mean square deviation (RMSD) with a 2.5 Å radius. The respective representation of chain C is shown in Fig. 4j.

**Supplementary Fig. 8.**
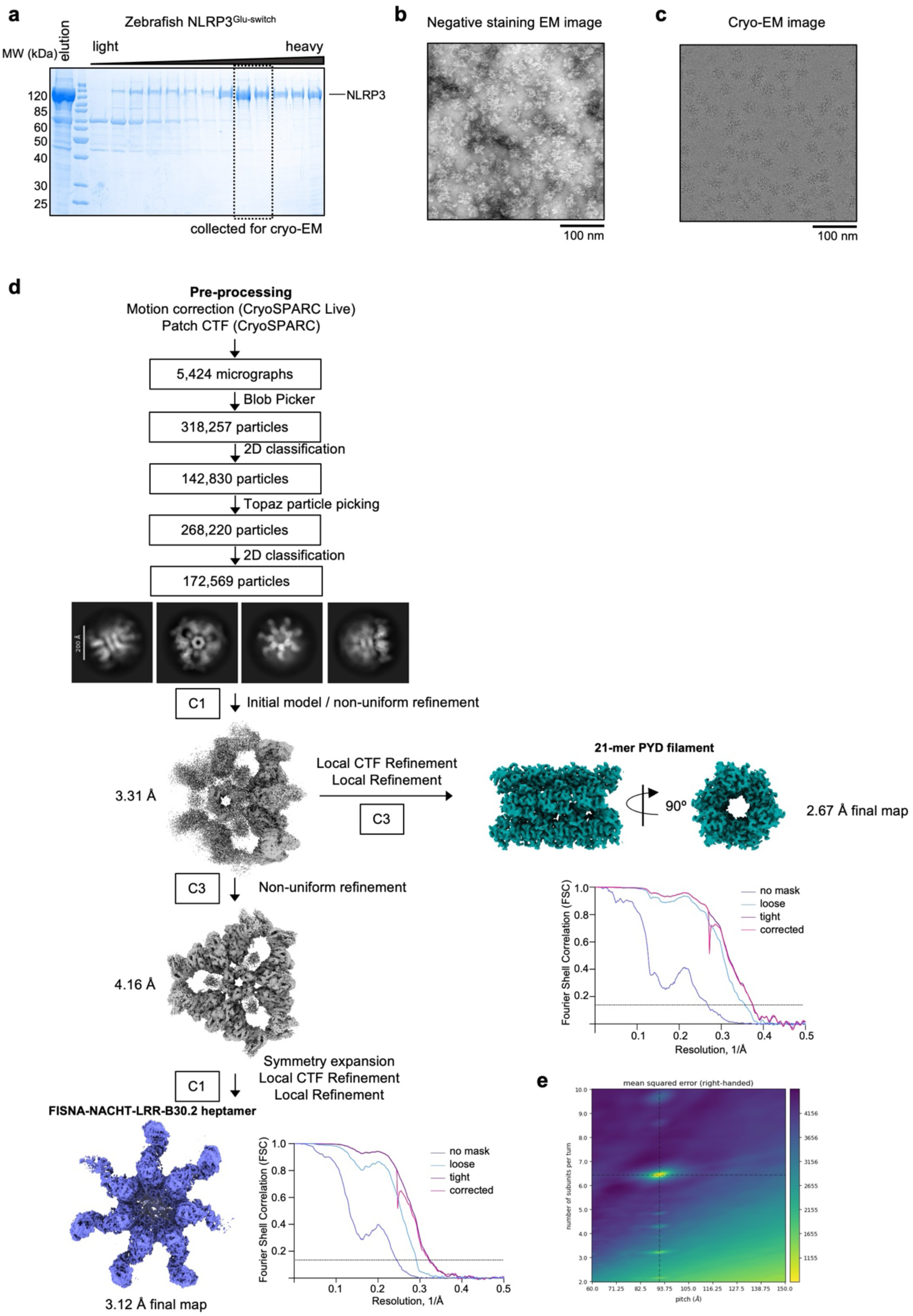
Structure determination workflow for the “Glu-switch”-mutated zebrafish NLRP3. **a**, SDS-PAGE gel of sucrose gradient fractions of the “Glu-switch”-mutated zebrafish NLRP3 (NLRP3^Glu-switch^). Fractions collected for cryo-EM are indicated with a dashed box. **b**, A representative negative-staining EM image of the zebrafish NLRP3 oligomer (scale bar 50 nm). **c**, A representative cryo-EM micrograph. **d**, Structure determination workflow with the cryo-EM map for the full “Glu-switch”-mutated zebrafish NLRP3 complex and final cryo-EM maps of a FISNA-NACHT-LRR-B30.2 domains-containing heptamer and PYD 21-mer, and corresponding Fourier Shell Correlation (FSC) curves. The dashed line indicates the 0.143 cutoff. **e**, 2D plot of the mean squared error surface along the helical pitch and number of subunits per turn. The best estimate of helical symmetry parameters is located on the interception of the global minima depicted with dashed lines. See also Supplementary Table 1.

**Supplementary Fig. 9.**
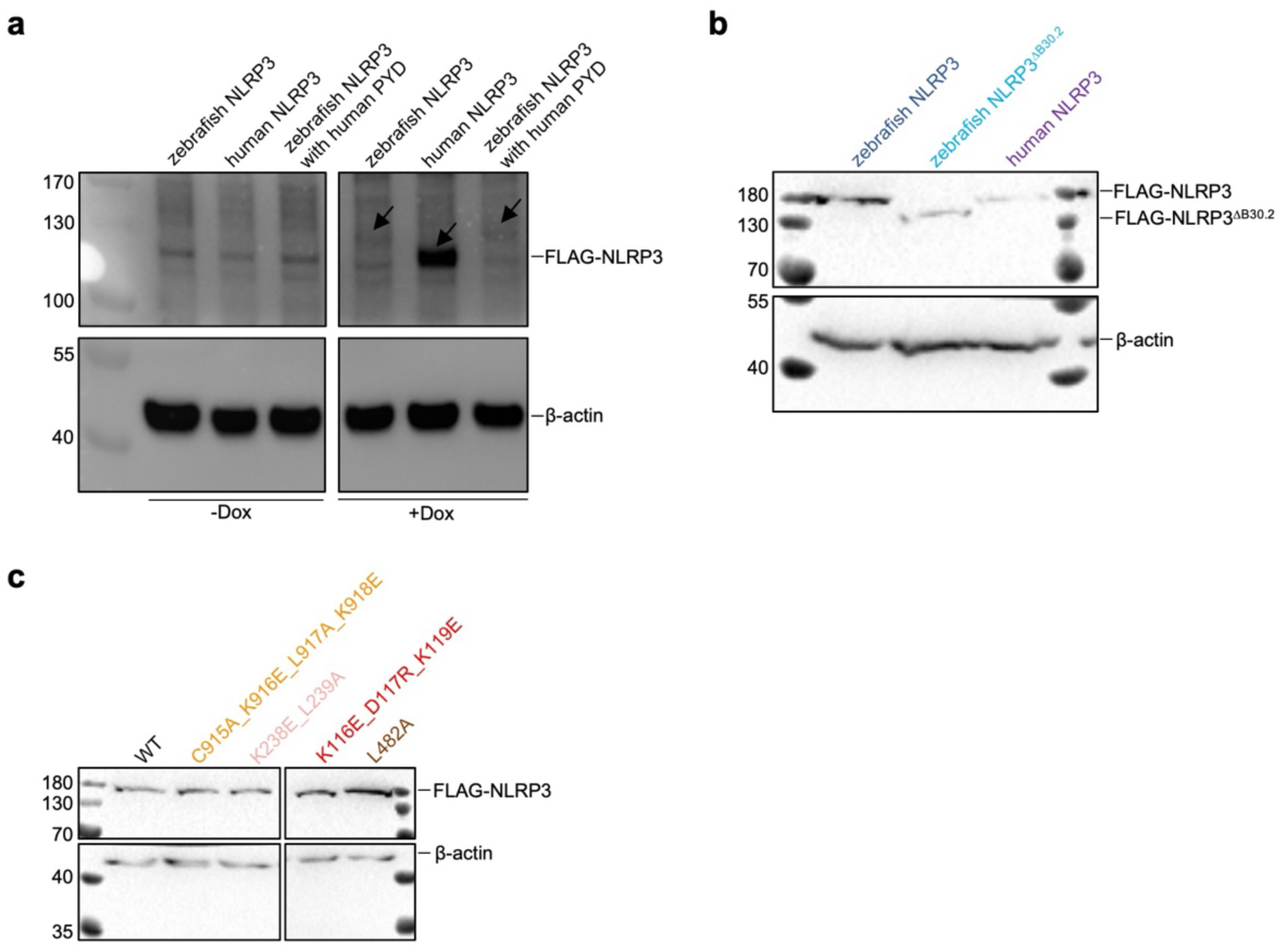
Expression levels of NLRP3 constructs in human cells. **a**, Western blot analysis of NLRP3 expression in reconstituted cell lines in Fig. 5a following doxycycline (Dox) induction. Bands corresponding to FLAG-tagged NLRP3 constructs are indicated with arrows. **b, c**, Western blot analysis of the whole cell lysates from HEK293T cells reconstituted with YFP-ASC and transfected with the indicated NLRP3 constructs, demonstrating relative expression levels of NLRP3 constructs used for Fig. 5c (**b**) and Fig. 5d (**c**). FLAG-tagged NLRP3 constructs and β-actin were visualized with corresponding antibodies. Cells were transfected with 100 ng DNA per well in a 96-well plate, and the lysates were collected 24 h post-transfection.

**Supplementary Fig. 10.**
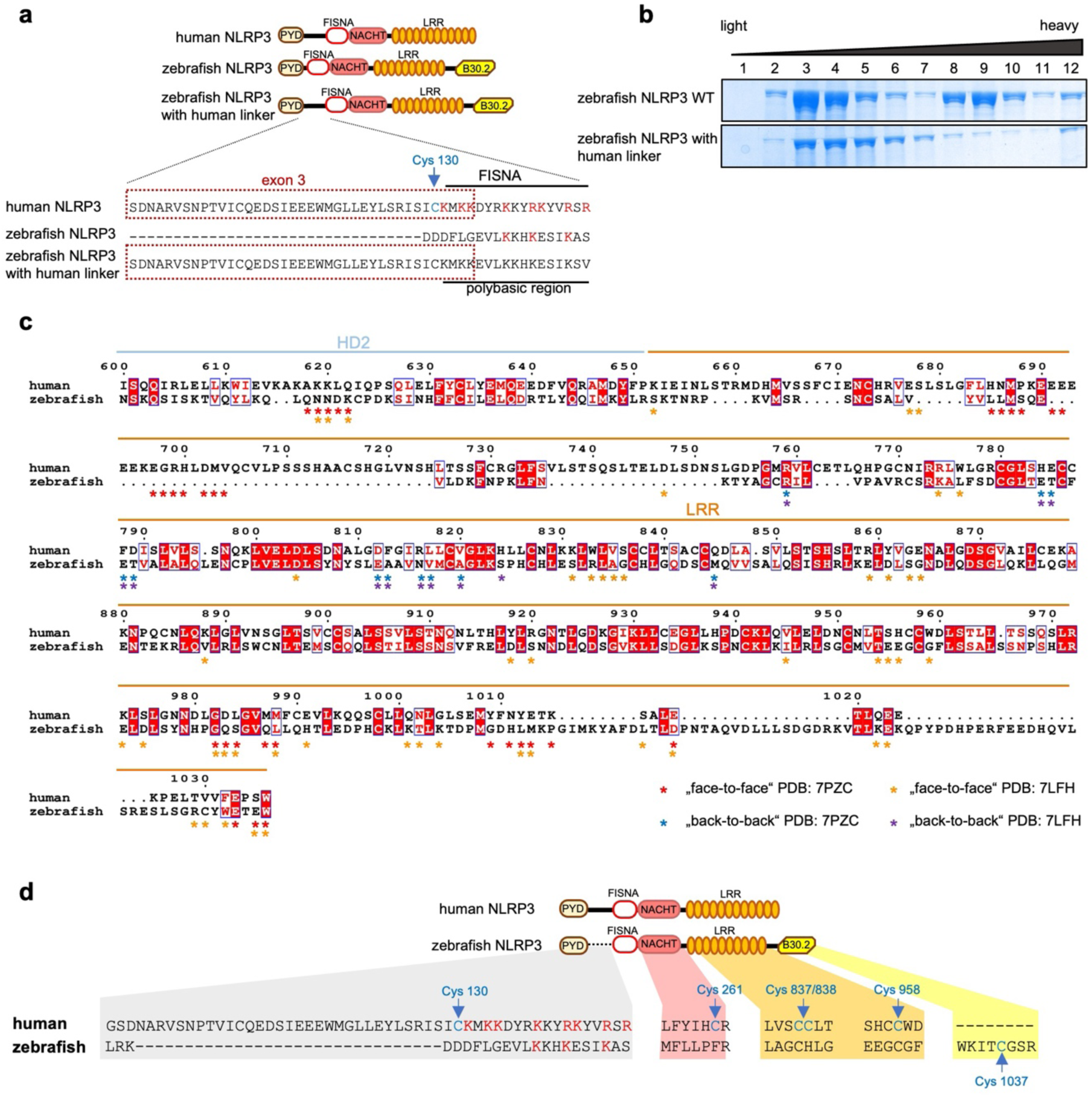
Comparison of the “cage”-specific sequences and palmitoylation sites in zebrafish and human NLRP3. **a**, Position and sequence of the PYD-FISNA linker in human and zebrafish NLRP3, and zebrafish NLRP3 containing human PYD-FISNA linker sequence encoded by the exon 3 of human *NLRP3* gene (red dashed box). Sequences corresponding to FISNA domain and polybasic region are indicated with black lines. Cys130 palmitoylated in human NLRP3 is highlighted in blue. **b**, SDS-PAGE gels of sucrose gradient fractions of zebrafish NLRP3 WT and zebrafish NLRP3 containing human linker sequence. **c**, Sequence alignment of zebrafish and human NLRP3. Domains are indicated with lines and color-coded. Residues forming a “face-to-face” interface according to the human (PDB: 7PZC) and mouse (PDB: 7LFH) NLRP3 “cage” structures are indicated with red and orange stars, respectively. Residues forming a “back-to-back” interface according to the human (PDB: 7PZC) and mouse (PDB: 7LFH) NLRP3 “cage” structures are indicated with blue and violet stars, respectively. Numbers correspond to alignment positions within the full-length sequence alignment. **d**, Position and sequence of known palmitoylation sites in human and zebrafish NLRP3. Palmitoylated residues are highlighted in blue.

**Supplementary Fig. 11.**
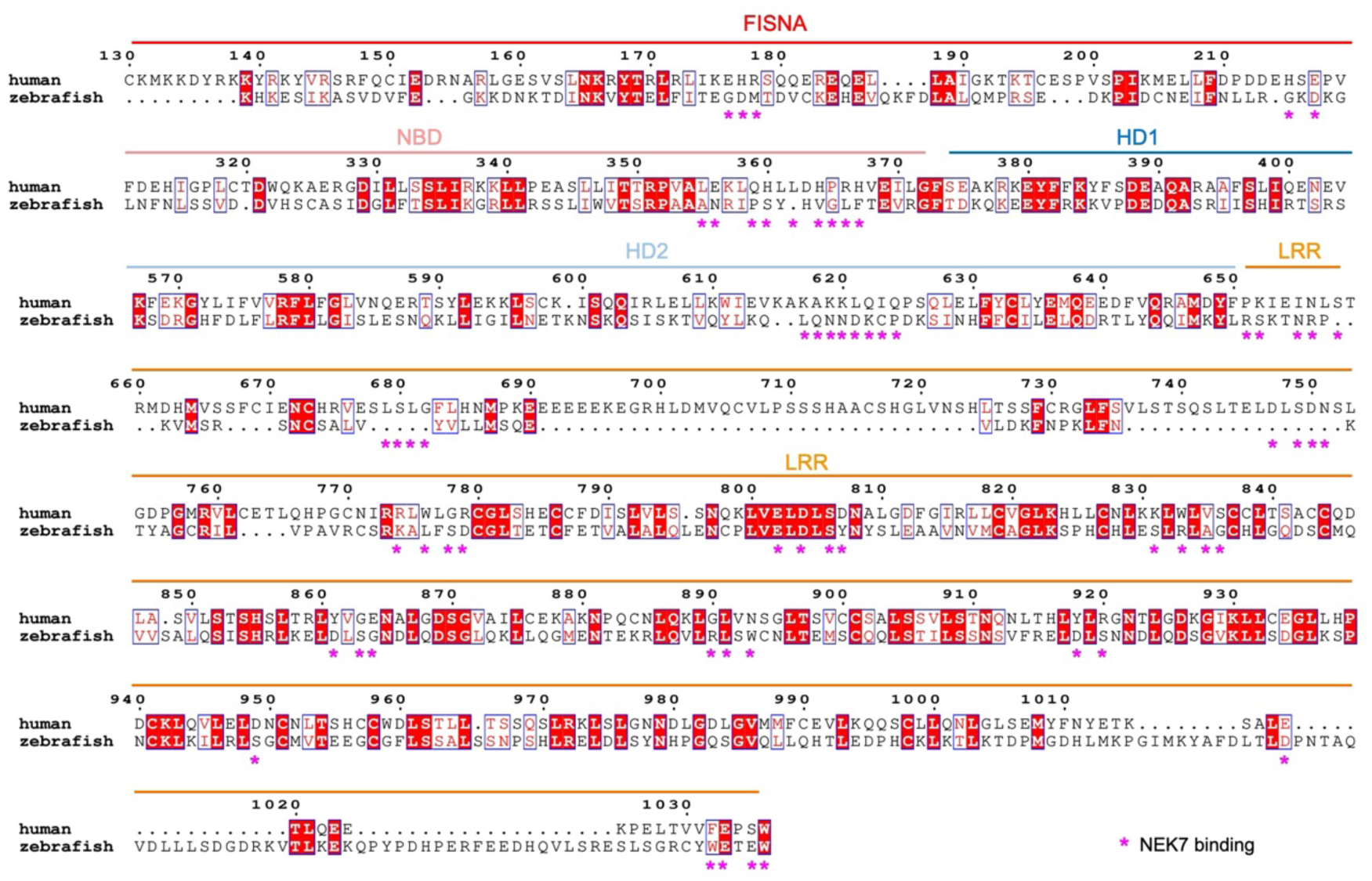
Zebrafish NLRP3 does not interact with NEK7. Sequence alignment of zebrafish and human NLRP3. Domains are indicated with lines and color-coded. Residues forming a NEK7-binding interface according to the human NLRP3:NEK7 complex structure (PDB: 6NPY) were identified using a PDBePISA^46^ analysis of human NLRP3:NEK7 complex structure^45^ and are indicated with magenta. NLRP3 sequence of 6NPY has been corrected using PDB: 7PZC as a reference.

## Data availability

The atomic coordinates and the corresponding cryo-EM density maps have been deposited in the Protein Data Bank and the Electron Microscopy Data Bank with the accession codes: PDB 29WX, EMD-57420 (wild-type zebrafish NLRP3 FISNA-NACHT-LRR-B30.2 heptamer), PDB 29WY, EMD-57421 (wild-type zebrafish NLRP3 PYD 21-mer), PDB 29WZ, EMD-57422 (“Glu-switch”-mutated zebrafish NLRP3 FISNA-NACHT-LRR-B30.2 heptamer), PDB 29XA, EMD-57423 (“Glu-switch”-mutated zebrafish NLRP3 PYD 21-mer). Any additional data are available from the corresponding author upon request.

## Acknowledgements

We gratefully acknowledge Jule Riebelmann and Anneli Yilmaz from the Core Facility Electron Microscopy at the University Hospital Tübingen for support, the electron microscopy facility at the Max Planck Institute for Biology Tübingen, especially Dr. Katharina Hipp, for screening, preliminary dataset collection, support and assistance in this work, Dr. Stefan Steimle, Birte Knickmeyer and Julia Seifermann for cryo-EM data collection, support and advice. We also thank Dr. Kristin Bieber from the Flow Cytometry Core Facility Tübingen Berg for cell sorting and analysis and Dr. Max Zimmermann from the Werner Siemens Imaging Center, Department of Preclinical Imaging and Radiopharmacy, University Hospital Tübingen for hardware and software support. Electron microscopy data were collected at the Cryo-EM Facility of the University of Freiburg (RRID: SCR_025860). The Titan Krios G4 cryo-TEM used for imaging was funded by Deutsche Forschungsgemeinschaft (project-ID 506518771) and is operated within the Microscopy and Image Analysis Platform (MIAP), University of Freiburg. pLenti-CMVie-IRES-BlastR (Addgene 119863) was a gift from Dr. Ghassan Mouneimne, pCMV4-HA (Addgene 27553) was a gift from Dr. Shao-Cong Sun, pET28-MBP-TEV (Addgene 69929) was a gift from Zita Balklava and Thomas Wassmer, lentivirus packaging plasmids pMD2.G (Addgene 12259) and psPAX2 (Addgene 12260) were gifts from Dr. Didier Trono, HEK293 Flp-In T-REx NLRP3-SH + ASC-EGFP cells were kindly provided by Prof. Dr. Florian I. Schmidt (Institute of Innate Immunity, University Hospital Bonn, Biomedical Center, Bonn, Germany). This work was supported by the DFG (German Research Foundation) Emmy Noether Programme (project-ID 520465503 to L.A.), DFG clusters of excellence “iFIT- Image Guided and Functionally Instructed Tumor Therapies” (EXC-2180 to A.N.R.W. and L.A.) and “CMFI- Controlling Microbes to Fight Infections” (EXC-2124 to A.N.R.W.), the Damon Runyon Cancer Research Foundation and CRIS Cancer Foundation (DFS-57-23 to L.A.), the University of Tübingen Medical Faculty (Fortüne-Antrag Nr. 2615-0-0 and Nr. 3023-0-0 to M.M.-T.), the ERC (ERC-2020-ADG–101018672 ENGINES, to V.H.) and the DFG (CRC1403, project-ID 414786233 to V.H.).

## Author contributions

Conceptualization: L.A., L.S.D.; Methodology: L.A., L.S.D., M.M.-T., V.V., V.H.; Investigation: L.S.D., C.G.-K., M.M.-T., V.V., M.H.W., L.S.; Validation: L.S.D, L.A.; Resources: L.S.D., C.G.-K., V.V., A.N.R.W., V.H., L.A.; Visualization: L.S.D., M.M.-T., V.V., L.A.; Writing-original draft: L.S.D., L.A.; Writing-review & editing: L.S.D., C.G.-K., M.M.-T., V.V., M.H.W., L.S., A.N.R.W., V.H., L.A.; Project administration and supervision: L.A.; Funding acquisition: L.A.

## Methods

### Constructs and cloning

Codon-optimized full-length (NCBI MN088121.1) and B30.2-deleted (NLRP3^ΔB30.2^, aa 1-923) zebrafish NLRP3, zebrafish NLRP3 with human linker (aa 95-134), human full-length and exon 3-deleted (aa 95-134) NLRP3 were cloned into pLenti CMVie-IRES-BlastR (Addgene, 119863) and modified by addition of an N-terminal FLAG-tag, mScarlet fluorescent protein and a Tobacco Etch Virus (TEV) cleavage site. Point mutations were introduced into zebrafish NLRP3 construct using the Q5 High-Fidelity 2x Master Mix (NEB): “Glu-switch” - R219E&E220R&L223A, interface I - C915A&K916E&L917A&K918E; interface II - K238E&L239A and N274A&L275A&S276A&S277A&D279R&R298E; interface III - W374A&T378A&V379A&D382R, L482A and E484R; interface IV - P318A&S319A&Y320C and K116E&D117R&K119E. Construct containing FLAG- and MBP-tagged human PYD-deleted NLRP3 (NLRP3 ^ΔPYD^, aa 126-1033) was described before^1^. HA-tagged full-length zebrafish (NCBI NP_001003617.1) and human NEK7 were cloned into pcDNA3.1 (Thermo Fisher) and pCMV-HA (Addgene, 27553), respectively. Zebrafish ASC^PYD^ (aa 1-89) with a mutation G89C was cloned into pET28-MBP-TEV (Addgene, 69929). Human full-length ASC was cloned into pLenti-EF1a-C-tGFP (Origene, PS100072) and tGFP was exchanged with YFP. For generation of stable BLaER1 cell lines, NLRP3 constructs were subcloned into a pLIX-Blast vector for doxycycline-inducible expression using Gibson assembly and an N-terminal FLAG tag was added.

### Protein expression and purification

For protein expression, one liter of Expi293F cells with a density of 3×10^6^ cells/ml were transfected with 1 mg plasmid using 1 mg polyethyleneimine (Polysciences, 23966-1) as transfection reagent. 72 h post-transfection cells were harvested by centrifugation. Cells were lysed by sonication (2 s on, 8 s off, 10 min total on, 40% power, Fisherbrand, 505) in buffer containing 50 mM Tris, 150 mM NaCl, 10% Glycerol, 5 mM MgCl_2_, pH7.5, supplemented with protease inhibitor (Sigma-Aldrich, 11836170001) and 2 mM β-mercaptoethanol. The lysate was clarified by centrifugation at 34,500 rcf and 4 °C for 1 h and used for affinity chromatography with anti-FLAG M2 Affinity Gel (Sigma-Aldrich, A2220). The protein was eluted with 100 μg/ml 3X FLAG-peptide (Biotrend, 331-11222-1) in a buffer containing 20 mM Tris, 150 mM NaCl, 2.5 mM MgCl_2_, pH 7.5 and incubated with 0.5 mM ATP for 30 min on ice. For Fig. 4c and Supplementary Fig. 7b elution was incubated with no nucleotide, 0.5 mM ADP or 0.5 mM AMP-PNP for 30 min on ice. The mixture was loaded onto a step-gradient of 20%, 30%, 35%, 40%, 45%, 50% and 60% sucrose in 20 mM Tris, 150 mM NaCl, 2.5 mM MgCl_2_, pH 7.5, supplemented with 0.2 mM of the respective nucleotide and centrifuged for 17 h at 136,000 rcf and 4 °C. The sucrose gradient was manually fractionated into 14 fractions and the zebrafish NLRP3-containing fractions were buffer exchanged using Zeba spin desalting columns (Thermo Fisher, A57763) equilibrated with buffer containing 20 mM Tris, 150 mM NaCl, 2.5 mM MgCl_2_, pH 7.5, supplemented with 0.2 mM of the same nucleotide as used for the sucrose gradient.

His_6_-MBP-tagged zebrafish ASC^PYD^ was expressed in *E. coli* strain BL21 (DE3) at 18°C overnight following the induction with 0.1 mM IPTG. Cells were lysed by sonication (3 s on, 5 s off, 5 min total on, 45% power, Fisherbrand, 505) in lysis buffer containing 20 mM Tris, 200 mM NaCl, 10% Glycerol, 5 mM MgCl_2_, pH 8.0, supplemented with 1 mM phenylmethanesulfonyl fluoride (Carl Roth, 6367.2) and 2 mM β-mercaptoethanol. The lysate was clarified by centrifugation at 34,500 rcf and 4 °C for 1 h and used for affinity chromatography using pre-equilibrated Ni-NTA Agarose (Quiagen, 30230) in a gravity-flow chromatography column (BioRad, 7321010). The protein was eluted in lysis buffer supplemented with 300 mM imidazole (Carl Roth, 3899.4) and further purified by size-exclusion chromatography using a Superdex 75 10/300 GL column (Cytiva) equilibrated with buffer containing 25 mM Tris, 150 mM NaCl, pH 8.0.

### Negative-staining electron microscopy

A 4 µl drop of sample was applied to a glow-discharged copper grid (Science Services, EFCF400H-Cu-50), incubated for 1 min, washed twice with buffer containing 20 mM Tris, 150 mM NaCl, 2.5 mM MgCl_2_, pH7.5, stained with 1% uranyl acetate for 40 s and air dried. The images were collected at 40,000x magnification at the Electron Microscopy Facility of the University Dermatology Clinic using Zeiss LIBRA 120 transmission electron microscope (Carl Zeiss) operated at 120 kV and at the Electron Microscopy Facility of the Max Plack Institute for Biology Tübingen using 120kV Tecnai G2 Sprit BioTWIN equipped with a TVIPS F416 camera.

For Fig. 3e 1 µM wild-type zebrafish NLRP3 was mixed with 10 µM of zebrafish ASC^PYD^ in a buffer containing 20 mM Tris, 150 mM NaCl, 2.5 mM MgCl_2_, 0.5 mM ATP, pH 7.5, and ASC^PYD^ polymerization was induced with 0.5 µM TEV at 4°C overnight. The complex was visualized as described above.

### Cryo-EM data collection

A 3 µl drop containing zebrafish NLRP3 complex was applied to a Quantifoil grid with ultrathin carbon support (Quantifoil R1.2/1.3 400 mesh Au + 2 nm C), incubated for 15 s, blotted for 3 s, plunged into liquid ethane, and flash frozen using a FEI Vitrobot Mark IV (Thermo Fisher) at 100% humidity and 4°C. Grid conditions were optimized using screening at the Electron Microscopy Facility of the Max Plack Institute for Biology Tübingen using 200kV FEI Talos Arctica microscope (Thermo Fisher) equipped with an autoloader and Falcon III direct electron detector. Final datasets were collected at the Cryo-EM Facility of the University of Freiburg (RRID: SCR_025860) using Krios G4 300 kV Transmission Electron Microscope (Thermo Fisher) equipped with an autoloader, Selectris energy filter and Falcon 4i direct electron detector. Automated data collection was performed using EPU software (Thermo Fisher). The movies were obtained in counting mode at 130,000x magnification (0.937 Å). All videos were recorded at multiple defocus values from -0.5 to -2.2 µm over 40 frames with a total dose 40 e^−^/Å^2^.

### Cryo-EM data processing

Raw movies were corrected by gain reference and beam-induced motion using CryoSPARC Live and used for further image processing using CryoSPARC^2^.

For the wild-type zebrafish NLRP3 dataset a total of 8,598 micrographs were used for initial particle picking applying the Blob Picker function in CryoSPARC^2^. Particles were extracted with a box size of 512 pixels followed by 2D classification. Selected 2D classes were used as templates for two rounds of Topaz automated particle picking^3^ followed by particle extraction and a 2D classification in CryoSPARC^2^, which resulted in a total of 91,512 particles in the selected 2D classes. Initial 3D models were generated by *ab initio* reconstruction^2^ and the best model was used for a homogenous refinement followed by a non-uniform refinement^4^ with no symmetry applied resulting in a 3.43 Å initial cryo-EM map. For PYD 21-mer reconstruction, local CTF and local masked refinement focusing on PYD 21-mer density and using C3 symmetry was performed resulting in a 2.62 Å final cryo-EM map. Helical symmetry parameters of the PYD 21-mer were estimated using the symmetry search tool in CryoSparc. For NLRP3 heptamer reconstruction, the initial cryo-EM map was subjected to a non-uniform refinement with a C3 symmetry^4^ and symmetry expansion. Local CTF and masked refinement focused on a single zebrafish NLRP3 heptamer resulted in a 3.18 Å final cryo-EM map.

A dataset of the “Glu-switch”-mutated zebrafish NLRP3 (R219E&E220R&L223A) contained 5424 micrographs and yielded 318,157 particles picked by the Blob Picker function in CryoSPARC^2^. Particles were extracted with a box size of 512 pixels and subjected to a 2D classification. Selected 2D classes were used as templates for the Topaz automated particle picking^3^ followed by particle subtraction and a 2D classification^2^, which resulted in total 172,569 particles in the selected 2D classes. Initial 3D models were generated with *ab initio* reconstruction^2^, and the best model was used for homogenous refinement followed by a non-uniform refinement^4^ with no symmetry applied resulting in a 3.31 Å initial cryo-EM map. For PYD 21-mer reconstruction, local CTF and local masked refinement focusing on PYD 21-mer density and using C3 symmetry was performed resulting in a 2.67 Å final cryo-EM map. Helical symmetry parameters of the PYD 21-mer were estimated using the symmetry search tool in CryoSPARC^2^. For NLRP3 heptamer reconstruction, the initial cryo-EM map was subjected to a non-uniform refinement with a C3 symmetry^4^ followed by a symmetry expansion. Local CTF and masked refinement focusing on a single NLRP3 heptamer resulted in a 3.12 Å final cryo-EM map.

Post-processing of all maps was performed with DeepEMhancer^5^ for model building and representation. Local resolution estimation for final maps was calculated with Local Resolution Estimation module in cryoSPARC^2^. Data processing statistics are summarized in the Supplementary Table 1.

### Model building and structure representation

Cryo-EM maps of the zebrafish NLRP3 heptameric discs were first fit with seven copies of the zebrafish NLRP3 model predicted by AlphaFold 3^6^ using UCSF Chimera^7^. First, FISNA-NBD-HD1 and WHD-HD2-LRR-B30.2 segments from the AlphaFold 3^6^ prediction were fitted separately using a rigid body fit followed by a rigid body fitting of the separate FISNA, NBD, HD1, WHD, HD2 and LRR-B30.2 domains. The model was further manually adjusted in Coot^8^ and subjected to a real-space refinement in Phenix^9^. Similarly, for PYD 21-mers reconstruction, 21 copies of zebrafish NLRP3^PYD^ (aa 1-90) predicted with AlphaFold 3^6^ were fitted into the cryo-EM maps of the PYD 21-mers using UCSF Chimera^7^ and subjected to a real-space refinement in Phenix^9^.

Interfaces were analyzed using PDBePISA^10^. Structure representations were made using UCSF Chimera^7^ and Pymol^11^.

### *In vitro* lipid blot assay

Lipid blot assay with purified NLRP3 protein constructs was performed using membrane lipid strips (Biozol, ECH-P-6002) as previously described^1^. In short, lipid strips were blocked with 3 % bovine serum albumin (BSA) in phosphate-buffered saline with 1% Tween 20 (PBS-T) for 1.5 h at room temperature (RT) followed by incubation with 21 µl FLAG-tagged NLRP3 proteins (13 µM) diluted in 3% BSA in PBS-T for 1 h at room temperature. The membrane strips were washed for a total of 20 minutes with PBS-T and the FLAG-tagged proteins were visualized using anti-FLAG-HRP antibodies (1:10,000, 4°C overnight, Sigma-Aldrich, A8592). All samples were visualized at the same time under the same conditions.

### Pull-down assay

FLAG-tagged NLRP3 and HA-tagged NEK7 constructs were co-expressed in 300 ml Expi293F cells transfected at the density 3×10^6^ cells/ml with 140 µg of each plasmid and 560 µg polyethyleneimine (Polysciences, 23966-1) as transfection reagent. 72 h post-transfection cells were harvested by centrifugation, lysed in buffer containing 50 mM Tris, 150 mM NaCl, 5 mM MgCl_2_, 10% Glycerol, pH 7.5, supplemented with protease inhibitor (Sigma-Aldrich, 11836170001) and 2 mM β-mercaptoethanol by sonication (2 s on, 8 s off, 2 min total on, 40% power, Fisherbrand, 505). The lysate was clarified by centrifugation at 34,500 rcf and 4 °C for 1 h and SDS samples were taken for loading controls. The remaining lysate was incubated for 1 h at 4°C with 300 µl anti-FLAG M2 Affinity Gel (Sigma-Aldrich, A2220) and washed with 10 ml of the buffer containing 50 mM Tris, 150 mM NaCl, 5 mM MgCl_2_, pH 7.5, followed by protein elution with 100 μg/ml 3X FLAG-peptide (Biotrend, 331-11222-1) in the same buffer. Lysate and elution samples were analyzed by Western blotting using anti-FLAG-HRP (1:10,000, Sigma-Aldrich, A8592) and anti-HA-HRP (1:2000, Abcam, ab173826).

### Cell culture and generation of stable cell lines

HEK293T were cultivated in Dulbecco’s Modified Eagle’s medium (DMEM, Thermo Fisher, 21969035), supplemented with 10% fetal bovine serum (Pan Biotech, P30-3306) and Antibiotika-Antimykotikum (Gibco, 15240062). THP-1 cells were cultivated in RPMI 1640 medium (Thermo Fisher, 11875093) supplemented with 10% fetal bovine serum (Pan Biotech, P30-3306) and Antibiotika-Antimykotikum (Gibco, 15240062). BLaER1 cells were cultivated in RPMI 1640 medium (Thermo Fisher) supplemented with 10% heat-inactivated fetal calf serum (Thermo Fisher), 100 U/mL penicillin/streptomycin (Thermo Fisher Scientific), and 1 mM sodium pyruvate (Thermo Fisher). All cells were cultivated at 37°C in 5% CO_2_.

Stable cell lines were generated using lentiviral transduction. For HEK293T and THP-1 stable line generation, 0.6×10^6^ HEK293T cells were transfected with 1 μg of the plasmid containing the gene of interest, 750 ng psPAX2 packaging plasmid (Addgene, 12260) and 250 ng pMD2.G envelope plasmid (Addgene, 12259) for lentivirus production. After 48 h the virus-containing medium was collected and filtered using a 0.45 μm filter (FischerScientific, 17104371). For HEK293T stable line generation the virus was applied to fresh HEK293T cells supplemented with 8 μg/ml polybrene (Santa-Cruz Biotechnology, sc-134220). For THP-1 stable line generation the NLRP3-KO THP-1 cells lines were resuspended in the lentivirus-containing medium supplemented with 8 μg/ml polybrene (Santa-Cruz Biotechnology, sc-134220) and centrifuged for 1.5 h at 1,000 rcf at RT. The supernatant was discharged and infected cells were incubated in fresh medium supplemented with 5 μg/ml blasticidin for selection. The cells were expanded and mScarlet- or YFP-positive cells were selected by cells sorting using a Sony MA900 multi-application cell sorter equipped with a 130 μm nozzle at 6-10 psi. HEK293T cells stably expressing full-length human ASC-YFP were sorted for low expression levels to avoid inflammasome-independent ASC speck formation.

HEK293 Flp-In T-REx NLRP3-SH + ASC-EGFP cells stably expressing human EGFP-tagged ASC were kindly provided by Prof. Dr. Florian I. Schmidt (Institute of Innate Immunity, University Hospital Bonn, Biomedical Center, Bonn, Germany)^12^.

For BLaER1 stable cell lines generation lentiviral particles were produced by reverse transfection of 3 × 10⁶ HEK293T cells with 1.5 µg pMDLg/pRRE, 1 µg VSV-G, 0.5 µg pRSV-REV, and 1 µg transfer plasmid using polyethylenimine “Max” (Polysciences, 24765) according to the manufacturer’s protocol. After 48 h, viral supernatants were harvested, centrifuged, and filtered through a 0.45 μm filter before transduction of BLaER1 cells. Successfully transduced cells were selected with 7 µg/mL blasticidin.

### ASC speck formation assay

6×10^5^ HEK293T cells stably expressing full-length human ASC-YFP were seeded into a 96-well plate. After 24 h cells were transfected with plasmids encoding mScarlet-tagged NLRP3 constructs using polyethyleneimine (Polysciences, 23966-1) as transfection reagent. DNA amounts per well are indicated in figure legends. 24 h post-transfection formation of the ASC specks was recorded in the green channel (excitation and emission wavelength ranges 461-487 nm and 500-530 nm, respectively) using Tecan Spark multimode plate reader (Tecan) equipped with a 10x objective. The number of ASC specks per image area were calculated using Fiji^13^ as the number of maxima in the green channel.

### Stimulation of inflammasome pathways

For transdifferentiation of BLaER1 monocytes into macrophages, 60,000 cells/well were plated into a 96-well plate and cultured in medium supplemented with 10 ng/ml M-CSF, 10 ng/ml IL-3, and 100 nM β-estradiol. After 5 days of differentiation, cells were used for stimulation experiments. To induce expression, 1 µg/ml doxycycline was added 18 h prior to stimulation. For NLRP3 activation, cells were primed with 200 ng/ml lipopolysaccharide (LPS, InvivoGen, tlrl-b5lps) for 4 h and subsequently stimulated with 6.5 µM nigericin (Sigma-Aldrich) or 30 µg/mL imiquimod (R837, InvivoGen) for 2 h. As a control, the NAIP–NLRC4 inflammasome was activated using 0.025 mg/ml anthrax toxin lethal factor fused to the Burkholderia T3SS needle protein (needle toxin, LFn-YscF), delivered together with 0.25 mg/ml protective antigen.

For NLRP3 activation in THP-1 cells, 1×10^6^ THP-1 NLRP3-KO cells reconstituted with NLRP3 constructs were seeded to a 6-well plate and transdifferentiated using 100 ng/ml Phorbol-12-myristat-13-acetate (PMA, Invivogen, tlrl-pma-2). After 24 h the medium was replaced and 24 h later the cells were used for stimulation experiments. For NLRP3 activation the cells were primed with 1 µg/ml LPS (Invivogen, tlrl-b5lps) for 3 h followed by NLRP3 activation with 20 µM nigericin (Tocris, 4312) or 200 µM imiquimod (Invivogen, tlrl-imq-10) for 1 h.

### LDH cytotoxicity assay

Supernatants from stimulated cells were collected and analyzed using the Pierce LDH Cytotoxicity Assay Kit (Thermo Fisher) to assess cell death. Results are presented relative to a lysis control from the same experiment, with values from doxycycline-untreated controls subtracted as background.

### Western blot analysi

BlaER1, THP-1 or HEK293T cells expressing inflammasome components were lysed in Laemmli buffer, and lysates were incubated at 95 °C for 5 min. FLAG-tagged NLRP3, β-actin and cleaved GSDMD were visualized by Western blotting using following antibodies: anti-FLAG M2-HRP (1:10,000, Sigma-Aldrich, A8592) and β-actin-HRP (1:1000, Santa Cruz Biotechnology, sc-47778) for BlaER1, anti-FLAG M2-HRP (1:10,000, Sigma-Aldrich, A8592), anti-β-actin (1:2000, Santa Cruz Biotechnology, sc-47778), anti-GSDMD (1:1000, Cell Signaling, 36425), anti-mouse-HRP (1:2,000, Cell Signaling, 7076S) and anti-rabbit-HRP (1:2,000, Cell Signaling, 7074S) for HEK293T and THP-1.

### Immunofluorescence (IF)

HEK293T cells were seeded on 12 mm coverslips at a density of 0,5 ×10^6^. For transiently transfected cell a transfection with 500 ng plasmid using Lipofectamine 2000 (Thermo Fisher) as transfection agent was performed 24 h prior to fixation. Cells were fixed with 4% paraformaldehyde (PFA) in PBS at 37 °C for 15 min, washed three times with 500 µl PBS and permeabilized with 0.05% saponin diluted in PBS for 5 min. Cells were then incubated in blocking buffer containing 2% BSA and 0.05% saponin in PBS for 1 h and incubated with primary antibodies for 2 h or overnight at 4°C. After incubation, cells were washed and incubated with secondary antibodies for 1 h at RT, washed with PBS and stained with Hoechst 33342 (1 µg/ml) for 10 min at RT. For negative controls the same staining was performed without primary antibodies.

Coverslips with the stained cells were mounted onto glass slides using the mounting media ProLong™ Diamond antifade mountant (Thermo Fisher) and the imaging was performed using a Zeiss LSM800 confocal microscope equipped with an Airyscan module and a Plan-Apochromat 63×/1.4 oil immersion objective. The system was controlled by Zeiss Zen Blue software, facilitating the acquisition and analysis of high-resolution images. For image acquisition, appropriate filter sets and sequential imaging were used to minimize bleed-through between fluorophores. Following antibodies were used for immunofluorescence: anti-RCAS1 (D2B6N, 1:100, Cell signaling, 12290S), anti-Pericentrin (1:500, Abcam, ab4448), Alexa Fluor™ 647 anti-Rabbit IgG (1:500, Invitrogen, A-21245).

### Quantification and statistical analysis

All experiments were performed as triplicates. For negative staining EM and immunofluorescence imaging representative images have been selected. NLRP3 bands in SDS-gels with sucrose gradient fractions were quantified using Fiji^13^ and displayed as sucrose gradient traces indicating a percentage of the NLRP3 protein in each fraction from the total amount of NLRP3 protein in the gradient. For all bar graphs, data were represented as mean ± SD of n=3 independent experiments with individual data points represented as circles.

---

**Supplementary Table 1.** Statistics of Zebrafish NLRP3 Structure Determination by Cryo-EM, Related to Figure 1 and Figure 4.

|  | **Zebrafish NLRP3 WT** |  | **Zebrafish NLRP3,**<br>**R219E_E220R_L223A** |  |
| --- | --- | --- | --- | --- |
| **Data collection and processing** | Heptamer disc | PYD filament | Heptamer disc | PYD filament |
| Magnification |  | 130,000 |  | 130,000 |
| Voltage (kV) |  | 300 |  | 300 |
| Electron exposure (e-/Å<sup>2</sup>) |  | ~40 |  | ~40 |
| Defocus range (μm) |  | -0.5 to -2.2 |  | -0.5 to -2.2 |
| Pixel size (Å) |  | 0.937 |  | 0.937 |
| Symmetry imposed | C1 | C3 | C1 | C3 |
| Initial particle images (no.) |  | 680,036 |  | 318,157 |
| Final particle images (no.) | 274,536 | 91,512 | 517,707 | 172,569 |
| Map resolution (Å) | 3.18 | 2.62 | 3.12 | 2.67 |
| FSC threshold | 0.143 | 0.143 | 0.143 | 0.143 |
| Local map resolution range (Å) | 2-10 | 2-11 | 2-15 | 2-8 |
| **Refinement** |  |  |  |  |
| Initial model used | AlphaFold 3 | AlphaFold 3 | AlphaFold 3 | AlphaFold 3 |
| Model resolution (Å) at FSC threshold | 1.8/ 2.1/ 3.5<br>0/ 0.143/ 0.5 | 1.6/ 1.8/ 2.7<br>0/ 0.143/ 0.5 | 1.9/ 2.1/ 3.5<br>0/ 0.143/ 0.5 | 1.6/ 1.8 / 2.7<br>0/ 0.143/ 0.5 |
| Map sharpening *B* factor (Å<sup>2</sup>) | -74.7 | -56.2 | -80 | -57.9 |
| **Model composition** |  |  |  |  |
| Non-hydrogen atoms | 57211 | 15309 | 57190 | 15309 |
| Protein residues | 7154 | 1890 | 7154 | 1890 |
| Ligands | 7 | 0 | 7 | 0 |
| ***B* factors (Å<sup>2</sup>)** |  |  |  |  |
| Protein | 123.32 | 61.35 | 123.7 | 62.93 |
| Ligand | 54.4 | - | 53.68 | - |
| **R.m.s. deviations** |  |  |  |  |
| Bond lengths (Å) | 0.003 | 0.004 | 0.003 | 0.003 |
| Bond angles (°) | 0.707 | 0.646 | 0.672 | 0.635 |
| **Validation** |  |  |  |  |
| MolProbity score | 2.28 | 1.87 | 2.29 | 1.99 |
| Clashscore | 11.24 | 8.15 | 10.6 | 9.15 |
| Poor rotamers (%) | 2.33 | 1.73 | 2.67 | 2.08 |
| **Ramachandran plot** |  |  |  |  |
| Favored (%) | 93.22 | 96.32 | 93.57 | 96.16 |
| Allowed (%) | 6.57 | 3.68 | 6.22 | 3.84 |
| Disallowed (%) | 0.21 | 0 | 0.21 | 0 |

## References

1. Putnam, C. D., Broderick, L. & HoAman, H. M. The discovery of NLRP3 and its function in cryopyrin-associated periodic syndromes and innate immunity. Immunol. Rev. 322, 259–282 (2024).

2. Swanson, K. V., Deng, M. & Ting, J. P.-Y. The NLRP3 inflammasome: molecular activation and regulation to therapeutics. Nat. Rev. Immunol. 19, 477–489 (2019).

3. Weber, A. N. R. et al. The expanding role of the NLRP3 inflammasome from periodic fevers to therapeutic targets. Nat. Immunol. 26, 1453–1466 (2025).

4. Coll, R. C. & Schroder, K. Inflammasome components as new therapeutic targets in inflammatory disease. Nat. Rev. Immunol. 25, 22–41 (2025).

5. Vande Walle, L. & Lamkanfi, M. Drugging the NLRP3 inflammasome: from signalling mechanisms to therapeutic targets. Nat. Rev. Drug Discov. 23, 43–66 (2024).

6. Bauernfeind, F. G., et al. Cutting Edge: NF-κB Activating Pattern Recognition and Cytokine Receptors License NLRP3 Inflammasome Activation by Regulating NLRP3 Expression. J. Immunol. 183, 787–791 (2009).

7. Dufies, O. & Zanoni, I. Post-translational modifications of NLRP3: to prime or not to prime? Trends Immunol. S1471490625002522 (2025) doi:10.1016/j.it.2025.10.008.

8. Barnett, K. C., Li, S., Liang, K. & Ting, J. P.-Y. A 360° view of the inflammasome: Mechanisms of activation, cell death, and diseases. Cell 186, 2288–2312 (2023).

9. Broz, P. & Dixit, V. M. Inflammasomes: mechanism of assembly, regulation and signalling. Nat. Rev. Immunol. 16, 407–420 (2016).

10. Martinon, F., Burns, K. & Tschopp, J. The Inflammasome. Mol. Cell 10, 417–426 (2002).

11. Andreeva, L. et al. NLRP3 cages revealed by full-length mouse NLRP3 structurecontrol pathway activation. Cell 184, 6299–6312.e22 (2021).

12. Hochheiser, I. V. et al. Structure of the NLRP3 decamer bound to the cytokine release inhibitor CRID3. Nature 604, 184–189 (2022).

13. Ohto, U., et al. Structural basis for the oligomerization-mediated regulation of NLRP3 inflammasome activation. Proc. Natl. Acad. Sci. 119, e2121353119 (2022).

14. Magupalli, V. G. et al. HDAC6 mediates an aggresome-like mechanism for NLRP3 and pyrin inflammasome activation. Science 369, eaas8995 (2020).

15. Xiao, L., Magupalli, V. G. & Wu, H. Cryo-EM structures of the active NLRP3 inflammasome disc. Nature 613, 595–600 (2023).

16. Glück, I. M. et al. Nanoscale organization of the endogenous ASC speck. iScience 26, 108382 (2023).

17. Masumoto, J. et al. ASC, a Novel 22-kDa Protein, Aggregates during Apoptosis of Human Promyelocytic Leukemia HL-60 Cells. J. Biol. Chem. 274, 33835–33838 (1999).

18. Netea, M. G. et al. DiAerential requirement for the activation of the inflammasome for processing and release of IL-1β in monocytes and macrophages. Blood 113, 2324–2335 (2009).

19. Gaidt, M. M. et al. Human Monocytes Engage an Alternative Inflammasome Pathway. Immunity 44, 833–846 (2016).

20. He, Y., Franchi, L. & Núñez, G. TLR agonists stimulate Nlrp3-dependent IL-1β production independently of the purinergic P2X7 receptor in dendritic cells and in vivo. J. Immunol. Baltim. Md 1950 190, 334–339 (2013).

21. Leal, V. N. C. et al. Bruton’s tyrosine kinase (BTK) and matrix metalloproteinase-9 (MMP-9) regulate NLRP3 inflammasome-dependent cytokine and neutrophil extracellular trap responses in primary neutrophils. J. Allergy Clin. Immunol. 155, 569–582 (2025).

22. Bork, F. et al. naRNA-LL37 composite DAMPs define sterile NETs as self-propagating drivers of inflammation. EMBO Rep. 25, 2914–2949 (2024).

23. Schmacke, N. A. et al. IKKβ primes inflammasome formation by recruiting NLRP3 to the trans-Golgi network. Immunity 55, 2271–2284.e7 (2022).

24. Wöhrle, S. et al. NEK7 accelerates NLRP3 inflammasome activation. Preprint at 10.1101/2025.09.26.678733 (2025).

25. Shi, H. et al. NLRP3 activation and mitosis are mutually exclusive events coordinated by NEK7, a new inflammasome component. Nat. Immunol. 17, 250–258 (2016).

26. Schmid-Burgk, J. L. et al. A Genome-wide CRISPR (Clustered Regularly Interspaced Short Palindromic Repeats) Screen Identifies NEK7 as an Essential Component of NLRP3 Inflammasome Activation. J. Biol. Chem. 291, 103–109 (2016).

27. He, Y., Zeng, M. Y., Yang, D., Motro, B. & Núñez, G. NEK7 is an essential mediator of NLRP3 activation downstream of potassium eAlux. Nature 530, 354–357 (2016).

28. Mateo-Tórtola, M. et al. Non-decameric NLRP3 forms an MTOC-independent inflammasome. Preprint at 10.1101/2023.07.07.548075 (2023).

29. Liu, Y. et al. Cryo-electron tomography of NLRP3-activated ASC complexes reveals organelle co-localization. Nat. Commun. 14, 7246 (2023).

30. Nie, L. et al. Consecutive palmitoylation and phosphorylation orchestrates NLRP3 membrane traAicking and inflammasome activation. Mol. Cell 84, 3336–3353.e7 (2024).

31. Yu, T. et al. NLRP3 Cys126 palmitoylation by ZDHHC7 promotes inflammasome activation. Cell Rep. 43, 114070 (2024).

32. Billman, Z. P., Hancks, D. C. & Miao, E. A. Unanticipated Loss of Inflammasomes in Birds. Genome Biol. Evol. 16, evae138 (2024).

33. Chen, H. et al. Characterization of the Japanese flounder NLRP3 inflammasome in restricting Edwardsiella piscicida colonization in vivo. Fish Shellfish Immunol. 103, 169–180 (2020).

34. Chen, S. et al. Dual function of a turbot inflammatory caspase in mediating both canonical and non-canonical inflammasome activation. Dev. Comp. Immunol. 121, 104078 (2021).

35. Li, N. et al. Palmitoylation-mediated NLRP3 inflammasome activation in teleosts highlights evolutionary divergence in immune regulation: *Zool*. Res. 46, 3–14 (2025).

36. Li, J.-Y. et al. The zebrafish NLRP3 inflammasome has functional roles in ASC-dependent interleukin-1β maturation and gasdermin E–mediated pyroptosis. J. Biol. Chem. 295, 1120–1141 (2020).

37. Tyrkalska, S. D. et al. Silica crystals activate toll-like receptors and inflammasomes to promote local and systemic immune responses in zebrafish. Dev. Comp. Immunol. 138, 104523 (2023).

38. Hu, Z. et al. Structural and biochemical basis for induced self-propagation of NLRC4. Science 350, 399–404 (2015).

39. Zhang, L. et al. Cryo-EM structure of the activated NAIP2-NLRC4 inflammasome reveals nucleated polymerization. Science 350, 404–409 (2015).

40. Matico, R. E. et al. Structural basis of the human NAIP/NLRC4 inflammasome assembly and pathogen sensing. Nat. Struct. Mol. Biol. 31, 82–91 (2024).

41. Acehan, D. et al. Three-Dimensional Structure of the Apoptosome. Mol. Cell 9, 423–432 (2002).

42. Yuan, S., Topf, M., Reubold, T. F., Eschenburg, S. & Akey, C. W. Changes in Apaf-1 Conformation That Drive Apoptosome Assembly. Biochemistry 52, 2319–2327 (2013).

43. Zhou, M. et al. Atomic structure of the apoptosome: mechanism of cytochrome c- and dATP-mediated activation of Apaf-1. Genes Dev. 29, 2349–2361 (2015).

44. Abramson, J. et al. Accurate structure prediction of biomolecular interactions with AlphaFold 3. Nature 630, 493–500 (2024).

45. Sharif, H. et al. Structural mechanism for NEK7-licensed activation of NLRP3 inflammasome. Nature 570, 338–343 (2019).

46. Krissinel, E. & Henrick, K. Inference of Macromolecular Assemblies from Crystalline State. J. Mol. Biol. 372, 774–797 (2007).

47. Hochheiser, I. V. et al. Directionality of PYD filament growth determined by the transition of NLRP3 nucleation seeds to ASC elongation. Sci. Adv. 8, eabn7583 (2022).

48. Danot, O., Marquenet, E., Vidal-Ingigliardi, D. & Richet, E. Wheel of Life, Wheel of Death: A Mechanistic Insight into Signaling by STAND Proteins. Structure 17, 172–182 (2009).

49. Brinkschulte, R. et al. ATP-binding and hydrolysis of human NLRP3. Commun. Biol. 5, 1176 (2022).

50. Duncan, J. A. et al. Cryopyrin/NALP3 binds ATP/dATP, is an ATPase, and requires ATP binding to mediate inflammatory signaling. Proc. Natl. Acad. Sci. 104, 8041–8046 (2007).

51. MacDonald, J. A., Wijekoon, C. P., Liao, K. & Muruve, D. A. Biochemical and structural aspects of the ATP-binding domain in inflammasome-forming human NLRP proteins. IUBMB Life 65, 851–862 (2013).

52. Leipe, D. D., Koonin, E. V. & Aravind, L. STAND, a Class of P-Loop NTPases Including Animal and Plant Regulators of Programmed Cell Death: Multiple, Complex Domain Architectures, Unusual Phyletic Patterns, and Evolution by Horizontal Gene Transfer. J. Mol. Biol. 343, 1–28 (2004).

53. Sandall, C. F., Ziehr, B. K. & MacDonald, J. A. ATP-Binding and Hydrolysis in Inflammasome Activation. Molecules 25, 4572 (2020).

54. Zhang, X. & Wigley, D. B. The ‘glutamate switch’ provides a link between ATPase activity and ligand binding in AAA+ proteins. Nat. Struct. Mol. Biol. 15, 1223–1227 (2008).

55. Aganna, E. et al. Association of mutations in the *NALP3/CIAS1/PYPAF1* gene with a broad phenotype including recurrent fever, cold sensitivity, sensorineural deafness, and AA amyloidosis. Arthritis Rheum. 46, 2445–2452 (2002).

56. Dodé, C. et al. New Mutations of CIAS1 That Are Responsible for Muckle-Wells Syndrome and Familial Cold Urticaria: A Novel Mutation Underlies Both Syndromes. Am. J. Hum. Genet. 70, 1498–1506 (2002).

57. Molina-López, C. et al. Pathogenic NLRP3 mutants form constitutively active inflammasomes resulting in immune-metabolic limitation of IL-1β production. Nat. Commun. 15, 1096 (2024).

58. Cosson, C. et al. Functional diversity of *NLRP3* gain-of-function mutants associated with CAPS autoinflammation. J. Exp. Med. 221, e20231200 (2024).

59. Rapino, F. et al. C/EBPα Induces Highly EAicient Macrophage TransdiAerentiation of B Lymphoma and Leukemia Cell Lines and Impairs Their Tumorigenicity. Cell Rep. 3, 1153–1163 (2013).

60. Coombs, J. R. et al. NLRP12 interacts with NLRP3 to block the activation of the human NLRP3 inflammasome. Sci. Signal. 17, eabg8145 (2024).

61. The Inflammasome: Methods and Protocols. vol. 1040 (Humana Press, Totowa, NJ, 2013).

62. Nizami, S. et al. A phenotypic high-content, high-throughput screen identifies inhibitors of NLRP3 inflammasome activation. Sci. Rep. 11, 15319 (2021).

63. Chen, J. & Chen, Z. J. PtdIns4P on dispersed trans-Golgi network mediates NLRP3 inflammasome activation. Nature 564, 71–76 (2018).

64. Cheng, T. C., Hong, C., Akey, I. V., Yuan, S. & Akey, C. W. A near atomic structure of the active human apoptosome. eLife 5, e17755 (2016).

65. Li, Y. et al. Mechanistic insights into caspase-9 activation by the structure of the apoptosome holoenzyme. Proc. Natl. Acad. Sci. 114, 1542–1547 (2017).

66. Yu, X. et al. Structural basis for the oligomerization-facilitated NLRP3 activation. Nat. Commun. 15, 1164 (2024).

67. Fu, J., Schroder, K. & Wu, H. Mechanistic insights from inflammasome structures. Nat. Rev. Immunol. 24, 518–535 (2024).

68. Boršić, E. et al. Clustering of NLRP3 induced by membrane or protein scaAolds promotes inflammasome assembly. Nat. Commun. 16, 4887 (2025).

69. Bittner, Z. A. et al. BTK operates a phospho-tyrosine switch to regulate NLRP3 inflammasome activity. J. Exp. Med. 218, e20201656 (2021).

70. Abramson, J. et al. Accurate structure prediction of biomolecular interactions with AlphaFold 3. Nature 630, 493–500 (2024).

## References

1. Andreeva, L. et al. NLRP3 cages revealed by full-length mouse NLRP3 structure control pathway activation. Cell 184, 6299–6312.e22 (2021).

2. Punjani, A., Rubinstein, J. L., Fleet, D. J. & Brubaker, M. A. cryoSPARC: algorithms for rapid unsupervised cryo-EM structure determination. Nat Methods 14, 290–296 (2017).

3. Bepler, T. et al. Positive-unlabeled convolutional neural networks for particle picking in cryo-electron micrographs. Nat Methods 16, 1153–1160 (2019).

4. Punjani, A., Zhang, H. & Fleet, D. J. Non-uniform refinement: adaptive regularization improves single-particle cryo-EM reconstruction. Nat Methods 17, 1214–1221 (2020).

5. Sanchez-Garcia, R. et al. DeepEMhancer: a deep learning solution for cryo-EM volume post-processing. Commun Biol 4, 874 (2021).

6. Abramson, J. et al. Accurate structure prediction of biomolecular interactions with AlphaFold 3. Nature 630, 493–500 (2024).

7. Meng, E. C. et al. UCSF ChimeraX: Tools for structure building and analysis. Protein Science 32, e4792 (2023).

8. Emsley, P., Lohkamp, B., Scott, W. G. & Cowtan, K. Features and development of *Coot*. Acta Crystallogr D Biol Crystallogr 66, 486–501 (2010).

9. Liebschner, D. et al. Macromolecular structure determination using X-rays, neutrons and electrons: recent developments in Phenix. Acta Crystallogr D Struct Biol 75, 861–877 (2019).

10. Krissinel, E. & Henrick, K. Inference of Macromolecular Assemblies from Crystalline State. Journal of Molecular Biology 372, 774–797 (2007).

11. The PyMOL Molecular Graphics System, Version 3.0, Schrödinger, LLC.

12. Tesfamariam, Y. M. et al. Poxvirus dsDNA genomes diAerentially activate AIM2 or NLRP3 inflammasomes in human primary cells. EMBO J 45, 1728–1759 (2026).

13. Schindelin, J., et al. Fiji: an open-source platform for biological-image analysis. Nat Methods 9, 676–682 (2012).

